# Sex-divergent sleep–wake network dysfunction after traumatic brain injury despite comparable tissue and neuronal loss

**DOI:** 10.64898/2026.11.20.768185

**Authors:** Audrey M. Magsig, Yuliya Voskobiynyk, Agnieszka Ciesielska, Drew Willoughby, Frances S. Cho, Jeanne T. Paz

## Abstract

The development of persistent sleep disruption, a common consequence after traumatic brain injury (TBI), is more likely to occur in female patients with TBI. Whether this difference stems from distinct structural tissue damage or functional network vulnerability remains unknown. Using continuous wireless electrocorticography in male and female mice, we evaluated sleep– wake network dynamics 3–4 weeks after focal cortical TBI (subacute-to-early-chronic phase). Females exhibited state-specific disruptions in delta (2–4 Hz) and mid-gamma (60–80 Hz) power—rhythms critical for sleep depth and arousal— whereas males displayed a broadband power reduction across all states. Both sexes showed disrupted sleep spindle magnitude, peak frequency, and interhemispheric coordination. Notably, lesion volumes and thalamic neurodegeneration were equivalent between sexes, but spatial transcriptomics revealed sex-divergent neuronal and immune responses in the thalamus. Together, these findings demonstrate that post-TBI sleep–wake network dysfunction is decoupled from gross structural tissue damage, establishing that sex-dependent network dynamics and distinct molecular responses shape functional recovery after TBI.

## INTRODUCTION

Traumatic brain injury (TBI) affects an estimated 50–70 million people worldwide each year and is a major cause of persistent neurological disability (*1–3*). Sleep–wake disturbances are among its most prevalent consequences, affecting 30% to 70% of patients (*4, 5*). These symptoms frequently persist throughout long-term recovery (*6*) and represent a major target for clinical intervention (*7*). Importantly, post-TBI sleep outcomes differ between females and males: large clinical studies associate female gender with more severe, persistent sleep disturbances after TBI (*6, 8*). However, whether these clinical disparities reflect biological, sex-dependent mechanisms or socio-environmental factors remains unclear.

Preclinical studies of post-traumatic electrophysiological sleep architecture have historically used male rodents (*9–13*). Consequently, how biological sex shapes persistent, state-dependent network oscillations and sleep microarchitecture, whether female estrous cycles modulate network stability, and how these functional phenotypes relate to underlying neurodegenerative pathology remain key open questions. Addressing these questions has critical implications for both experimental modeling and clinical trial design.

Following TBI, neurodegenerative processes do not remain static; rather, primary focal cortical injury triggers progressive, delayed pathology across connected subcortical networks— including the thalamus—over weeks to months (*9, 14, 15*). Preclinical work establishes that sleep–wake disturbances evolve during this subacute 2–8 week “latent” window, coinciding with secondary neurodegeneration and circuit remodeling in connected hubs (*9, 10, 12, 16*). This subacute period represents a critical window of circuit plasticity during which functional network vulnerabilities manifest. However, whether biological sex modulates subcortical network vulnerability and whether functional network disruptions reflect structural neurodegeneration during this critical window remains unknown.

Here, we investigated how biological sex influences sleep–wake outcomes 3 weeks post-injury in a mouse model of focal cortical TBI. We used continuous wireless electrocorticography (ECoG) to track sleep–wake architecture and state-dependent network activity in male and freely cycling female CD-1 mice subjected to controlled cortical impact (CCI) and examined whether post-injury phenotypes varied across the female estrous cycle. We further assessed whether sex-divergent functional outcomes were accompanied by differences in primary lesion burden, secondary neuronal loss, and neuroinflammation, and used spatial transcriptomics to characterize sex-dependent injury responses. We found distinct ECoG responses in females and males despite comparable changes to sleep–wake macroarchitecture, gross lesion size, and thalamic neuronal loss, together with sex-divergent molecular responses in the thalamus. These findings establish biological sex as a dynamic modifier of post-traumatic network vulnerability that acts independently of gross structural damage. Incorporating sex biology into investigations of post-traumatic sleep outcomes is essential for designing new TBI-specific therapies.

## RESULTS

### Electrocorticographic spectral responses to TBI differ between females and males across sleep–wake states

To determine how biological sex influences sleep–wake activity after TBI, we performed continuous wireless ECoG recordings in male and freely cycling female mice for 5 days beginning 3 weeks after CCI (*9*). Control animals underwent sham surgery without craniotomy. Recordings were classified as Active Wake (low delta, mid gamma power, high activity rate), Quiet Wake/REM (low delta, high theta/delta ratio, low activity rate), or NREM (high delta, low gamma power, low theta/delta ratio; Fig. 1B; see Methods). Estrous stage was determined by daily vaginal cytology in females, with equivalent handling in males.

**Fig. 1.**
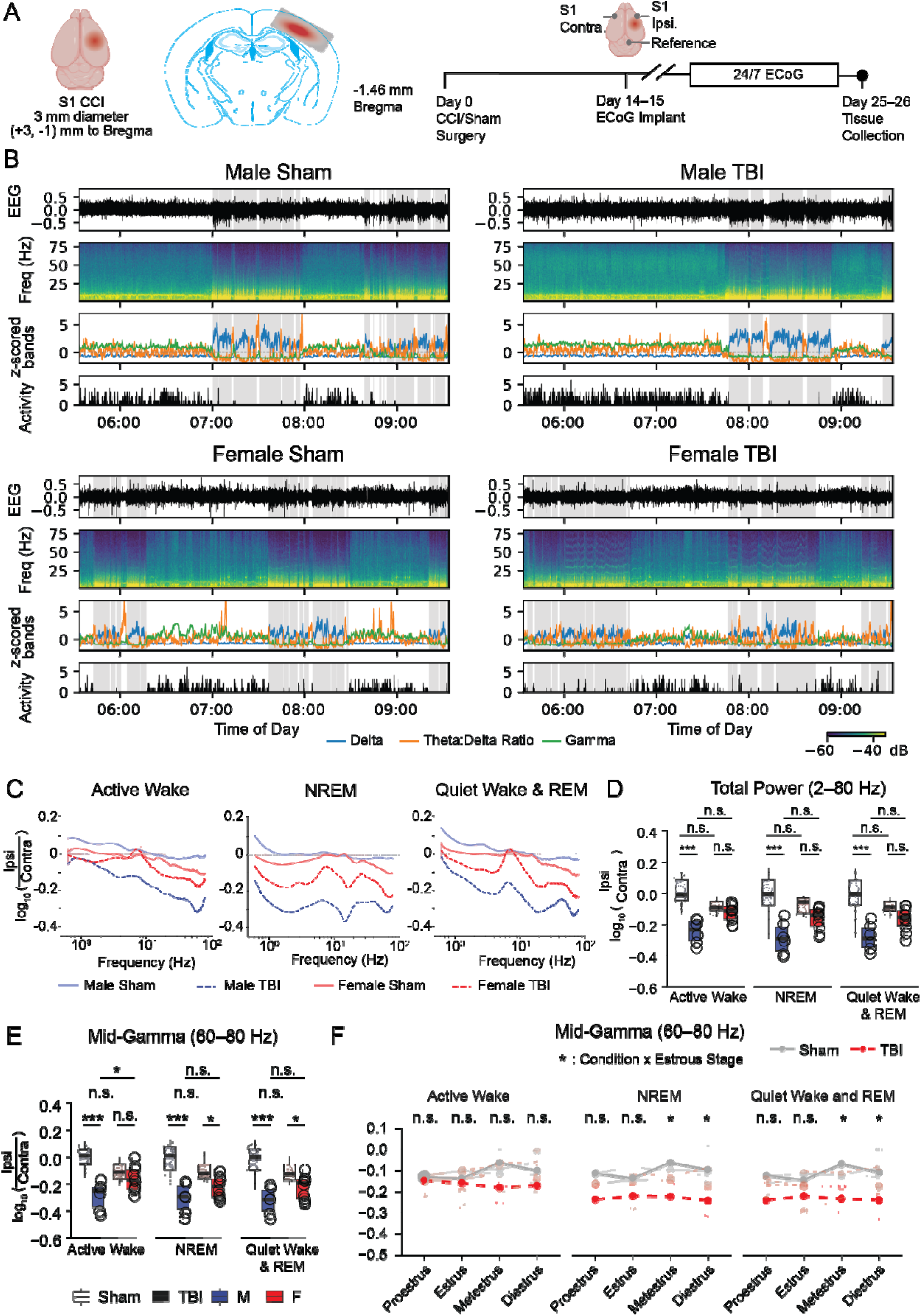
TBI broadly reduces ECoG spectral power in males but produces frequency-and state-specific reductions in females. (A) Left: Dorsal and coronal views of controlled cortical impact (CCI) targeting the S1 cortex. Right: Experimental timeline and ECoG electrode placement over ipsilateral (ipsi) and contralateral (contra) S1. (B) Representative 4-hour recording segments. From top to bottom: raw ECoG trace, spectrogram, z-scored features used for Hidden Markov model (HMM)-based sleep–wake classification, and activity counts. The horizontal dashed line indicates the mean of each z-scored feature. Gray-shaded regions indicate NREM-classified segments; unshaded regions correspond to Wake/REM. (C) Average log□□-transformed ipsi-to-contra ratio of spectral power across vigilance states and averaged across mice. Shading represents the standard error of the mean (SEM). The horizontal dashed line at 0 indicates equal power between hemispheres. (D) Average log□□-transformed ipsi-to-contra ratio of total power within vigilance state. Each point represents the mean value for one animal across all recorded. (E) Same as in (D) for mid-gamma power. (F) Average log□□-transformed ipsi-to-contra ratio of mid-gamma power across the estrous cycle in female mice. Each point represents the mean value per animal during a given estrous stage, lines connect values from one mouse. Dark lines represent group means; shading represents the 95% confidence interval. Linear mixed-effects models were used for statistical analyses. Detailed multiple-comparison procedures for individual analyses are reported in Table S1. n.s., not significant; * adjusted *p* < 0.05; ** adjusted *p* < 0.01; *** adjusted *p* < 0.001. Number of mice: (C–E) Male sham, n = 10; male TBI, n = 6; female sham, n = 7; female TBI, n = 10. (F) Female shams: proestrus, n = 4; estrus, n = 6; metestrus, n = 5; diestrus, n = 7; female TBIs: proestrus, n = 6; estrus, n = 8; metestrus, n = 8; diestrus, n = 9.

We first evaluated overall sleep–wake macro-architecture. TBI produced minimal effects on gross sleep–wake state architecture, consistent with reports in this strain of mice (Fig. S1A– C)(*10*). Across animals, TBI was only associated with an overall reduction in Active Wake bout number and an increase in mean bout duration, although these differences were not detected within either sex when analyzed separately (Fig. S1A–C). In females, the effect of TBI on the percentage of time spent in Active Wake varied with estrous stage, whereas no estrous-dependent effects were detected for the other sleep–wake measures examined (Fig. S1G–I). Sleep–wake measures varied with Zeitgeber time (ZT), as expected, but we found no evidence that TBI effects were restricted to a particular time of day (Fig. S1J–L; Table S1).

Because altered spectral power within canonical frequency bands is strongly linked to subjective sleep quality and circuit dysfunction (*17–19*), and gross sleep–wake architecture remained largely preserved, we next asked whether TBI altered the spectral composition of ECoG activity within individual sleep–wake states, whether these effects differed between females and males, and whether TBI effects in females varied across the estrous cycle. We quantified power in the delta (2–4 Hz), theta (6–9 Hz), sigma (10–15 Hz), beta (16–30 Hz), low-gamma (30–60 Hz), and mid-gamma (60–80 Hz) frequency bands and calculated the ratio of ipsilateral (injured hemisphere) to contralateral (uninjured hemisphere) ECoG power (Fig. 1A, B; see Methods for frequency-band selection).

In females, TBI reduced ipsilateral relative to contralateral mid-gamma power during NREM and Quiet Wake/REM, but not during Active Wake (Fig. 1C–E). This effect varied with estrous stage during NREM and Quiet Wake/REM, with significant differences between sham and TBI females during metestrus and diestrus (Fig. 1F). Similarly, the effect of TBI on low-gamma power varied with estrous stage (Fig. S3) although low-gamma power appeared unaltered by TBI when averaged across estrous stages (Fig. S2). In contrast, males exhibited reduced ipsilateral relative to contralateral power across nearly all frequency bands and sleep– wake states, consistent with previous observations (*9, 20*) (Fig. S2). Thus, TBI produced broad reductions in ECoG spectral power in males but frequency-and state-specific reductions in females, with some female spectral responses varying across the estrous cycle.

Because broad changes in spectral power can obscure frequency-specific differences between behavioral states, we next asked whether TBI altered relative spectral power between NREM and Active Wake. We calculated the ratio of NREM to Active Wake power within each ECoG electrode for each frequency band (Fig. 2A). At the level of total power, TBI did not significantly alter this ratio in either females or males (Fig. 2B). Frequency-specific analysis, however, revealed reduced NREM:Active Wake power ratios in the delta, theta, and mid-gamma bands in females, whereas reductions in males were restricted to the theta band (Fig. 2C). No detectable changes were observed in the other frequency bands or in the contralateral ECoG. Thus, TBI disrupted state-dependent theta activity exclusively in the injured hemisphere in both sexes, with additional disruptions in delta and mid-gamma activity in females.

**Fig. 2.**
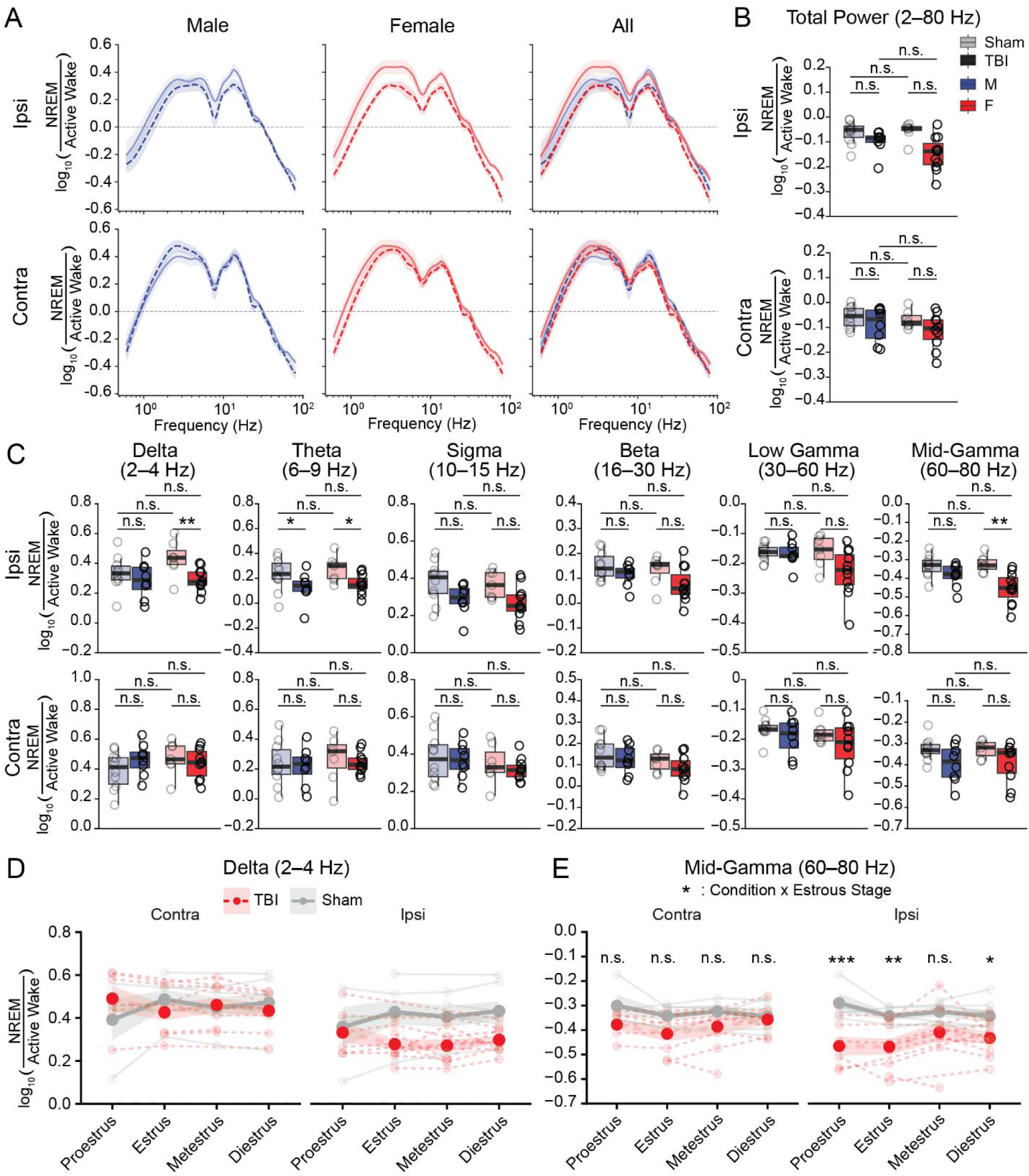
TBI produces distinct state-dependent spectral changes in females and males (A) Log□□-transformed ratio of average NREM power to average Active Wake power across all bouts for all considered frequencies. Traces represent group averages; shading represents SEM. Horizontal dotted line at 0 indicates equal power during NREM and Active Wake. (B) Log□□-transformed ratio of NREM to Active Wake total power. Each point represents one mouse. (C) Same as (B) for delta, theta, sigma, beta, low-gamma, and mid-gamma power. (D) Log□□-transformed ratio of NREM to Active Wake delta power across estrous stages in female mice. Each point represents the mean value per animal during a given estrous stage, lines connect values from one mouse. Light lines represent individual animals; dark lines represent group means; shading indicates 95% confidence interval. (E) Same as in (D) for mid-gamma power. Linear mixed-effects models were used for statistical analyses. Multiple-comparison procedures for individual analyses are reported in Table S1. n.s., not significant; * adjusted *p* < 0.05; ** adjusted *p* < 0.01; *** adjusted *p* < 0.001. Number of mice: (A–C) Male sham, n = 10; male TBI, n = 8; female sham, n = 7; female TBI, n = 11. (D–E) Female shams: proestrus, n = 4; estrus, n = 6; metestrus, n = 5; diestrus, n = 7; female TBIs: proestrus, n = 6; estrus, n = 8; metestrus, n = 8; diestrus, n = 9 for the contralateral ECoG and n = 10 for the ipsilateral ECoG.

We next examined whether these female spectral responses varied across the estrous cycle. The TBI-associated reduction in the delta NREM:Active Wake power ratio did not vary significantly across the estrous stage (Fig. 2D). Similarly, TBI effects did not vary with estrous stage across most frequency bands (Fig. S4). In contrast, the mid-gamma NREM:Active Wake power ratio showed a significant TBI × estrous-stage interaction (Fig. 2E). Post-hoc comparisons identified significant differences between sham and TBI females during proestrus, estrus, and diestrus, but not metestrus. Overall, TBI-associated changes in mid-gamma power were particularly sensitive to estrous stage.

### TBI disrupts sleep spindle properties and interhemispheric coordination in both sexes

Sleep spindles, rhythms generated by the thalamus during NREM sleep, are disrupted after TBI in male mice (*9*). We therefore asked whether TBI affected spindle properties differently in females and males. Spindles were detected during NREM sleep using Morlet wavelet convolution, and spindle density, magnitude, internal frequency, and duration were quantified (Fig. 3A; see Methods).

**Fig. 3.**
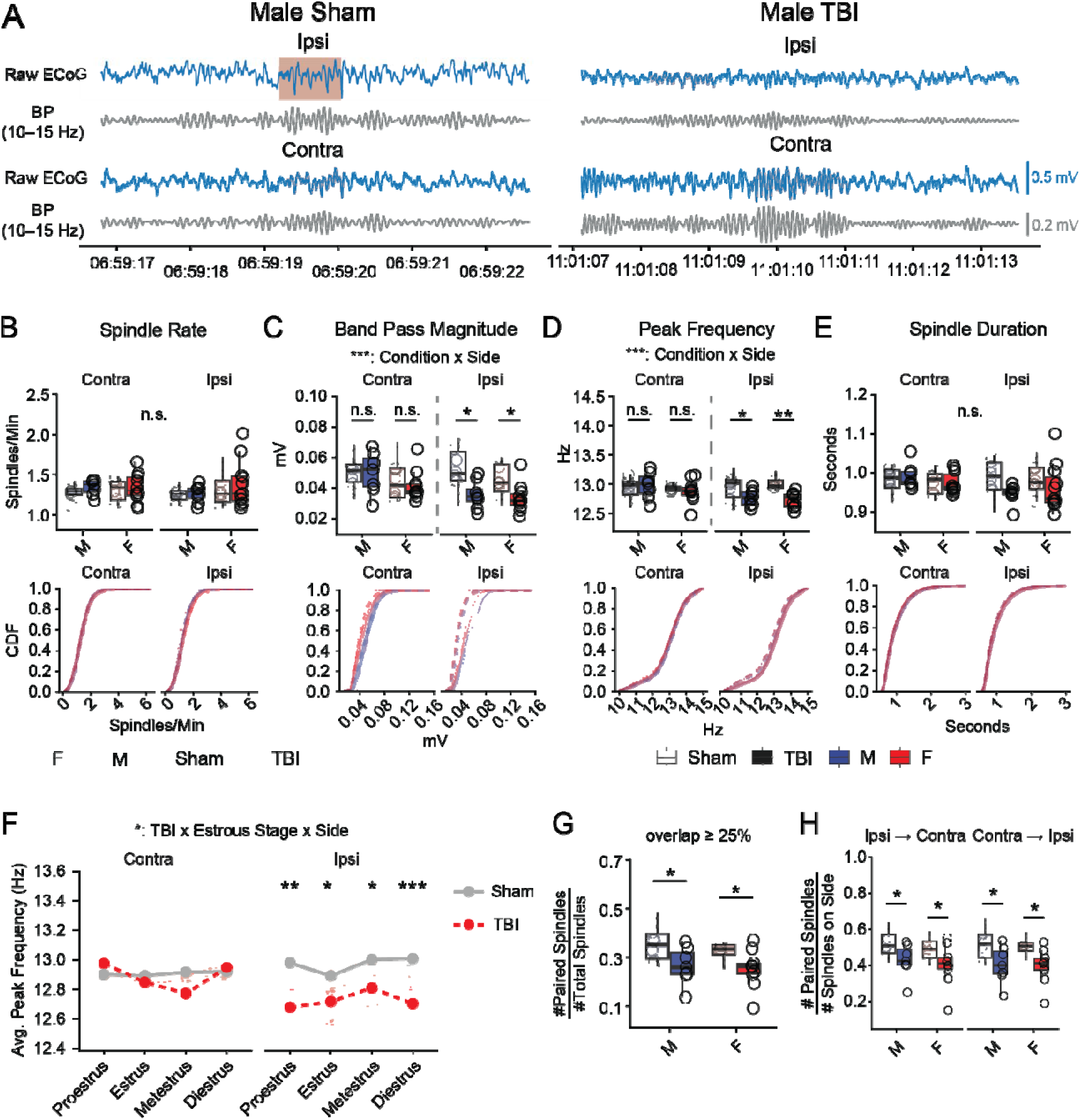
TBI disrupts sleep spindle properties and interhemispheric coordination in both sexes. (A) Representative sleep spindle detection. Shaded regions indicate detected spindles. (B) Top: Average spindle rate for each electrode. Bottom: Cumulative distribution functions (CDFs) for individual mice (light traces) and group average (dark traces). (C) Same as in (B) for the magnitude of the band-pass filtered signal. N = 1592–3647 spindles per animal. (D) Same as in (B) for peak frequency. (E) Same as in (B) for spindle duration. (F) Average spindle peak frequency across estrous stages in female mice. Proestrus: 44–79 spindles per mouse; estrus: 36–118; metestrus: 17–103; diestrus: 40–229. Dark lines show group means; shading indicates the 95% confidence interval; and light lines represent individual mice. (G) Proportion of unique spindles paired across hemispheres. (H) Same as in G split by hemispheric comparison. Each point represents one mouse. Linear mixed-effects models were used for statistical analyses. Multiple-comparison procedures for individual analyses are reported in Table S1. n.s., not significant; * adjusted *p* < 0.05; ** adjusted *p* < 0.01; *** adjusted *p* < 0.001. Number of mice: (A–E, G–H) male sham, n = 10; male TBI, n = 8; female sham, n = 8; female TBI, n = 11. (F): female shams: proestrus, n = 4; estrus, n = 6; metestrus, n = 5; diestrus, n = 8; female TBIs: proestrus, n = 5; estrus, n = 9; metestrus, n = 5; diestrus, n = 10.

Spindle magnitude and peak frequency were reduced in the ipsilateral ECoG of both sexes, whereas spindle rate and duration were unchanged (Fig. 3B–E). In females, the effect of TBI on spindle peak frequency also varied with estrous stage (Fig. 3F).

Because spindle timing is coordinated across brain regions (*21–23*), we next asked whether TBI disrupted spindle coordination across hemispheres. Spindles overlapping by at least 25% of their duration were classified as paired, and the ratio of paired to unpaired spindles was quantified across the recording. TBI reduced spindle pairing in both sexes, both overall (Fig. 3G) and when spindles originating from the ipsilateral or contralateral ECoG were considered separately (Fig. 3H). Thus, in contrast to the sex-divergent spectral responses across sleep–wake states, TBI-associated reductions in spindle magnitude, peak frequency, and interhemispheric coordination were largely shared between females and males.

### Cortical lesion volume and regional neuroinflammation are comparable between sexes

Previous studies have reported sex differences in edema and inflammatory responses within the primary injury site during the first week after TBI in animal models (*24–27*). We therefore asked whether the sex-divergent electrophysiological responses observed 3 weeks after TBI were accompanied by differences in cortical injury severity. Cortical lesion volume was assessed by profilometry (Fig. 4A)(*28*). Although TBI produced substantial cortical lesion size, lesion volume was comparable between males and females (Fig. 4B).

**Fig. 4.**
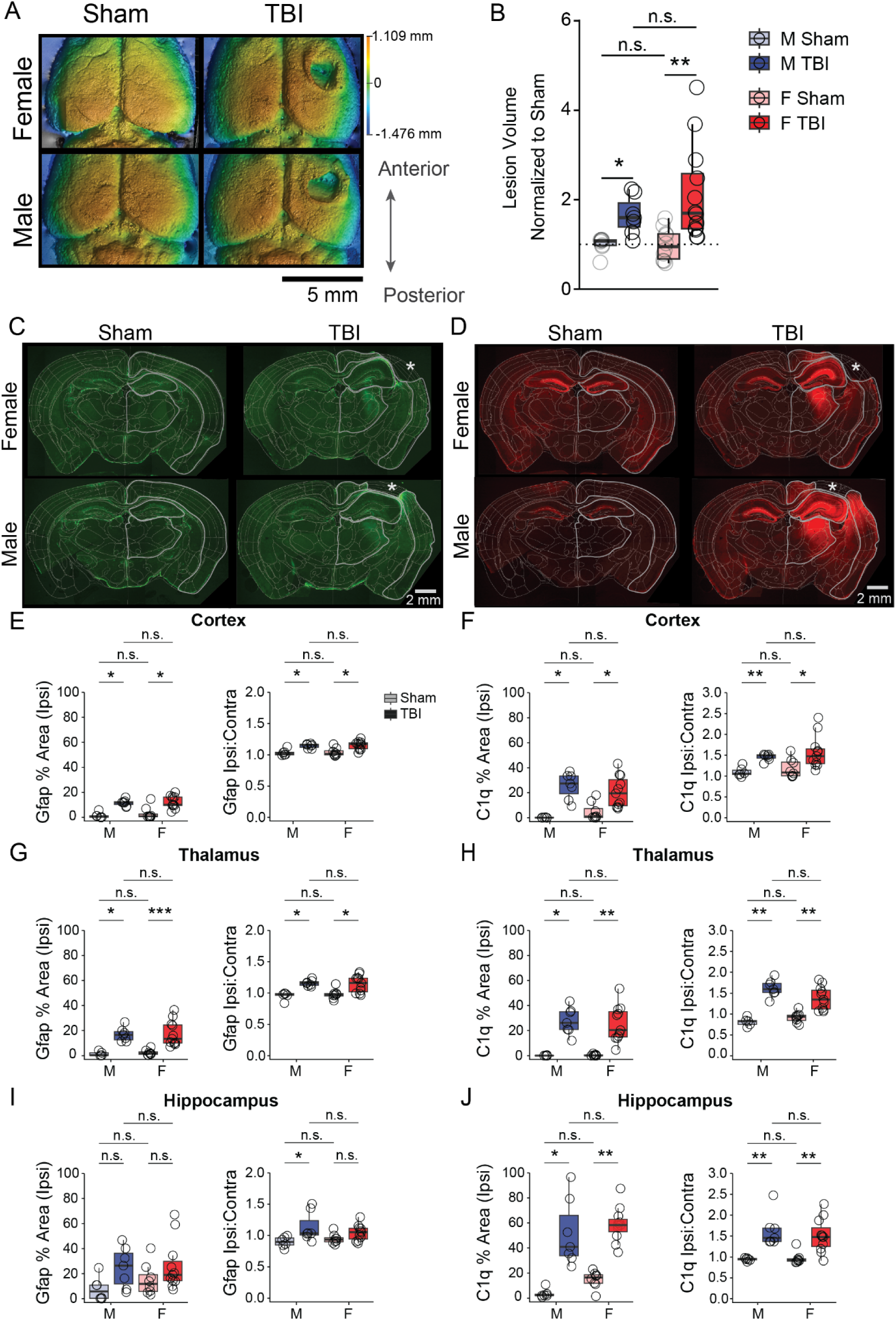
Cortical injury and regional neuroinflammation are broadly comparable between sexes after TBI. (A) Examples of surface profilometry images of the cortical surface 3 weeks after sham or TBI surgery. (B) Total displaced volume from the plane surrounding the TBI normalized to average sham values for the respective sex. The dotted line marks the sham average. (C) Representative images of GFAP staining in coronal sections from sham and TBI mice. Thin white lines indicate ROIs generated by registering sections to the Allen Brain Atlas using ABBA. Cortex, hippocampus, and thalamus are outlined in light gray. Asterisks indicate the cortical injury site. (D) Same as in (C) for C1q. (E) Cortex. Left: Percentage of the ipsilateral ROI covered by GFAP staining averaged per mouse. Right: Ipsilateral:contralateral GFAP fluorescence ratio. (F) Same as in (E) for C1q. (G–J) Same analyses as in (E, F) for the thalamus and hippocampus. Two-sided Wilcoxon rank-sum tests. Multiple-comparison procedures for individual analyses are reported in Table S1. n.s., not significant; * adjusted *p* < 0.05; ** adjusted *p* < 0.01; *** adjusted *p* < 0.001. Number of mice: (B) male sham, n = 8; male TBI, n = 7; female sham, n = 10; female TBI, n = 12. (E–J) male sham, n = 5; male TBI, n = 7; female sham, n = 8; female TBI, n = 11. Each data point represents the mean value for one mouse averaged across 2–4 sections from the indicated region.

We next examined whether regional inflammatory responses differed between sexes. Complement factor 1q (C1q), an upstream component of the classical complement pathway, and glial fibrillary acidic protein (GFAP), a marker of reactive astrocytes, were quantified in the peri-injury cortex, hippocampus, and thalamus and normalized to the respective region in the uninjured hemisphere. TBI increased C1q and GFAP in the cortex and thalamus to a similar extent in males and females (Fig. 4C–H). In the hippocampus, TBI increased C1q similarly in both sexes, but only increased GFAP in males (Fig. 4I, J). GFAP levels, however, did not differ significantly between male and female TBI animals. Thus, neither cortical lesion volume nor broad regional measures of neuroinflammation revealed substantial sex differences that paralleled the distinct electrophysiological responses to TBI.

Because secondary damage after TBI occurs in the specific subnuclei of the thalamus which are functionally connected to the injured cortex (*14*), we next used atlas-based brain registration with ABBA (Aligning Big Brains & Atlases) to systematically quantify C1q and GFAP across individual thalamic nuclei (Fig. 5A, B)(*29*). TBI increased both C1q and GFAP across sensory-associated nuclei, including the ventroposteromedial (VPM), ventroposterolateral (VPL), lateral dorsal (LD), posterior (PO), and reticular (RT) nuclei in both males and females (Fig. 5A–F). C1q increases also extended to the mediodorsal (MD), ventral anterior-lateral (VAL), and ventromedial (VM) nuclei, whereas GFAP increases were more restricted (Fig. 5G– J). Despite this widespread response to TBI, few sex differences were detected. C1q expression was greater in male than female TBI mice in the RT and MD nuclei, whereas GFAP did not differ between sexes in these regions. Thus, the regional inflammatory response to TBI was largely shared between sexes, with sex differences in C1q restricted to specific thalamic nuclei.

**Fig. 5.**
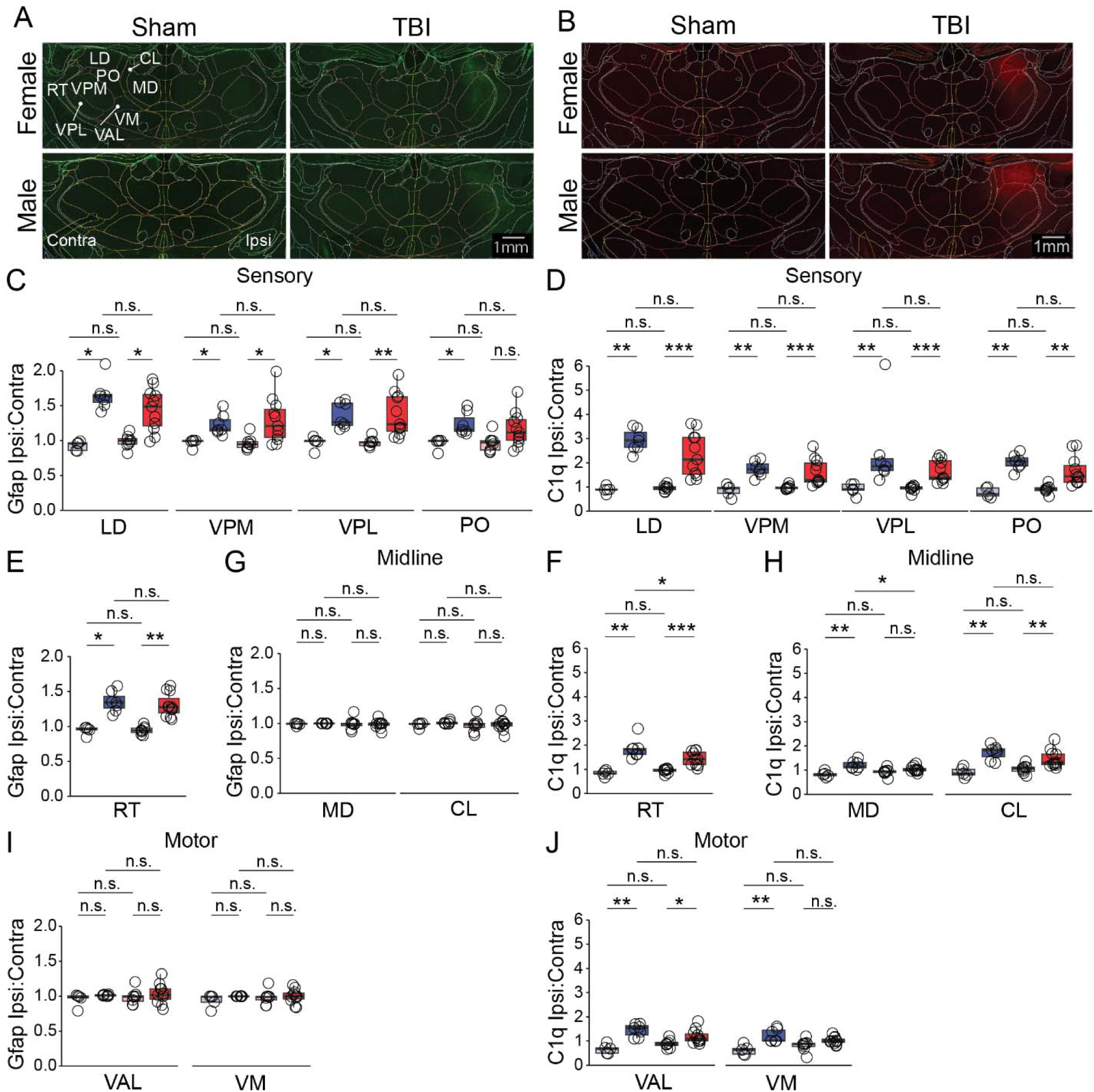
Thalamic inflammation is mainly biased towards sensory nuclei, with limited nucleus-specific sex differences in C1q expression. (A) Representative images of GFAP staining across thalamic nuclei in coronal sections from sham and TBI mice. Overlays indicate thalamic nuclei identified by registration to the Allen Brain Atlas, quantified nuclei are labeled in white, contra = contralateral/uninjured hemisphere, ipsi = ipsilateral/injured hemisphere. (B) Same as in (A) for C1q. (C) Ipsilateral:contralateral fluorescence ratios for GFAP in sensory-associated thalamic nuclei. (D) Same as in (C) for C1q. (E) Ipsilateral:contralateral fluorescence ratio for GFAP in the reticular thalamic nucleus (RT). (F) Same as in (E) for C1q. (G) Ipsilateral:contralateral fluorescence ratios for GFAP in midline thalamic nuclei. (H) Same as in (G) for C1q. (I) Ipsilateral:contralateral fluorescence ratios for GFAP in motor-associated thalamic nuclei. (J) Same as in (I) for C1q. Wilcoxon rank-sum tests, with FDR correction for comparisons within each stain. n.s., not significant; * adjusted *p* < 0.05; ** adjusted *p* < 0.01; *** adjusted *p* < 0.001. Raw and adjusted p values are reported in Table S1. Number of mice: (C–J) male sham, n = 5; male TBI, n = 7; female sham, n = 8; female TBI, n = 11. Each data point represents the mean value for one mouse averaged across 2–4 sections from the indicated region. Abbreviations for thalamic nuclei: CL, central lateral; LD, lateral dorsal; MD, mediodorsal; PO, posterior; RT, reticular thalamic; VAL, ventral anterior-lateral; VM, ventromedial; VPL, ventroposterolateral; VPM, ventroposteromedial.

We next used the same atlas-based approach to determine whether secondary thalamic neuronal loss or morphology differed between females and males. We quantified NeuN^+^ neurons and neuronal soma size within individual thalamic nuclei using an automated CellProfiler pipeline validated by manual analysis. Among sensory-associated nuclei, LD showed a substantial reduction in NeuN^+^ cells after TBI, but neuronal loss did not differ between sexes (Fig. 6A–C). Other sensory nuclei did not show detectable neuronal loss but exhibited reductions in neuronal soma size (Fig. 6D). Among TBI animals, neuronal soma size was smaller in females than males specifically in the VPL, with no sex differences detected in the other sensory nuclei examined (Fig. 6D). TBI did not alter neuronal soma size in the RT, and no sex differences were detected in any subregion (Fig. 6E). Within motor-associated nuclei, neuronal soma size was modestly reduced in the VAL of male TBI mice relative to male controls, but neither VAL nor VM differed between male and female TBI animals (Fig. 6F). Midline nuclei showed no detectable changes in neuronal soma size following TBI (Fig. 6G).

**Fig. 6.**
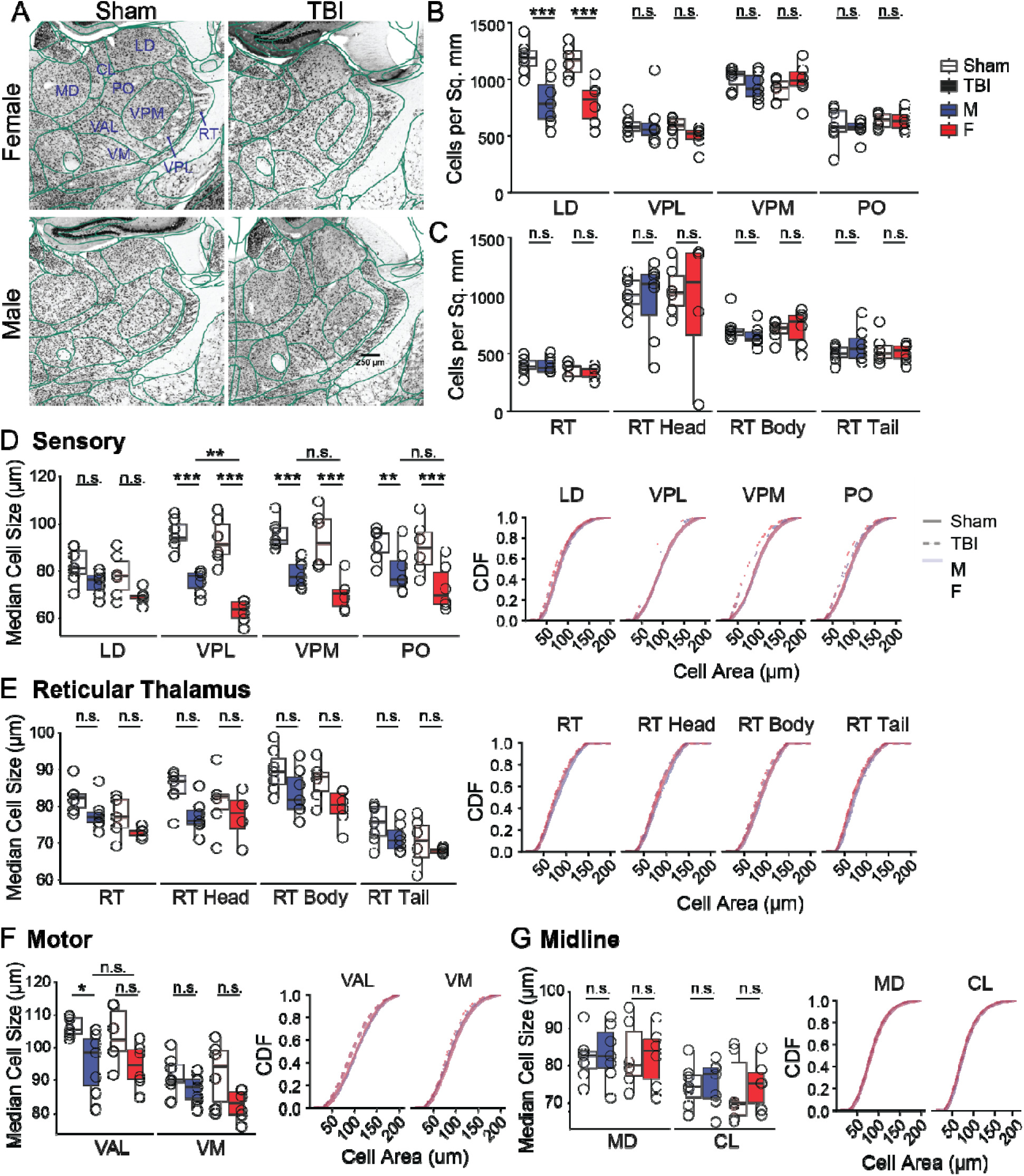
Secondary thalamic neuronal alterations are largely comparable between sexes. (A) Representative images of NeuN DAB staining in sham and TBI mice. Allen Brain Atlas registration is outlined in dark green. (B) NeuN^+^ cell density on the ipsilateral side across sensory-associated thalamocortical nuclei. (C) Same as in (B) for the RT and corresponding subregions. (D) Neuronal soma size across sensory-associated thalamic nuclei. Left: Median cell size for each mouse. Right: Cumulative distributions of cell area for male sham, male TBI, female sham, and female TBI mice; shading represents SEM across mice. (E) Same as in (D) for the RT and corresponding subregions. (F) Same as in (D) for motor-associated thalamic nuclei. (G) Same as in (D) for midline thalamic nuclei. Linear mixed-effects models were used for statistical analyses, with FDR correction across regions. Sex effects were tested only in regions showing a TBI effect and were FDR-corrected across all regions tested for sex effects. n.s., not significant; * adjusted *p* < 0.05; ** adjusted *p* < 0.01; *** adjusted *p* < 0.001. Detailed statistical results are reported in Table S1. Number of mice: (B–G) male sham, n = 7; male TBI, n = 7; female sham, n = 6; female TBI, n = 6. Each point represents the mean value for one mouse across 2–5 sections from the indicated region. For the RT head, two female TBI mice were excluded because the ROI could not be reliably quantified.

Together, these analyses show that cortical lesion size and secondary thalamic neuronal loss are broadly comparable between females and males 3 weeks after TBI. Inflammatory responses across the cortex, hippocampus, and thalamus were also largely shared between sexes, although localized differences in C1q expression and neuronal morphology were detected in specific thalamic nuclei.

### Spatial transcriptomics identifies sex-dependent responses in the thalamus after TBI

To determine whether males and females exhibit distinct responses in surviving thalamic tissue, we performed exploratory spatial transcriptomic profiling of somatosensory relay (VPL and VPM) and reticular thalamic (RT) regions using the NanoString GeoMx platform (Fig. 7A, B). Regions of interest (ROIs) were defined using NeuN and IBA1 immunoreactivity to enrich for neuronal and microglial compartments, respectively. Reliable segmentation of IBA1-enriched ROIs was not possible because of marked differences in IBA1 signal between sham and TBI tissue; analyses were therefore restricted to the whole somatosensory thalamic region and NeuN-enriched ROIs. Given the limited number of animals, differential expression analyses were exploratory, using uncorrected *p*-values to identify candidate sex-dependent responses for further study.

**Fig. 7.**
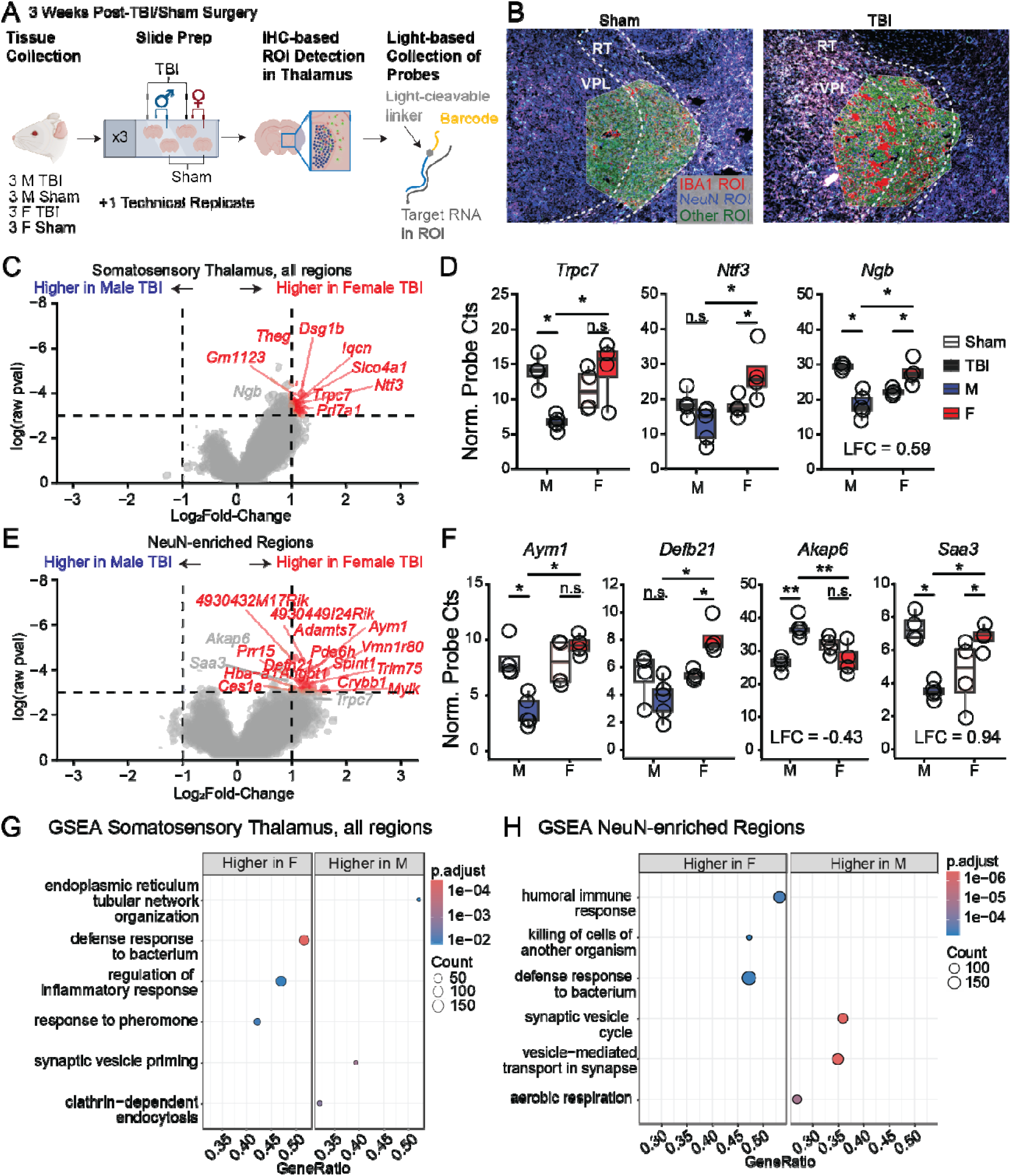
Spatial Transcriptomics suggests sex-divergent molecular programs in the thalamus after TBI. (A) Experimental design for spatial transcriptomics. ROIs were defined by immunohistochemistry-guided segmentation. (B) Representative segmentation of IBA1^+^, NeuN^+^, and IBA1^−^/NeuN^−^ ROIs in the approximate somatosensory regions of the VPL and RT (boundaries outlined in white). (C) Volcano plot comparing gene expression between female and male TBI mice across all ROIs. Horizontal dotted line indicates the exploratory threshold of uncorrected p = 0.05; Annotated genes differed between sham and TBI mice of either sex at *p* < 0.05 and |log□FC| >1 and were also altered by TBI relative to sham in at least one sex (*p* < 0.05), or were selected for (E, F). LFC, log□ fold change. (D) Representative genes illustrating sex-dependent expression patterns across all ROIs. Genes with |log□FC| < 1 are labeled with the corresponding effect size. (E) Same as in (C) for NeuN^+^ ROIs. (F) Same as in (D) for the NeuN^+^ ROIs. (G) Gene-set enrichment analysis of biological processes across all ROIs. (H) Same as in (G) for NeuN^+^ ROIs. Detailed statistical results are reported in Table S1. n.s., not significant; * *p* < 0.05; ** *p* < 0.01; *** *p* < 0.001. Panel A was created using BioRender.com. Number of mice: male sham, n = 3; male TBI, n = 3; female sham, n = 3; female TBI, n = 3. (C– H) Number of ROIs: male sham, n = 4; male TBI, n = 5; female sham, n = 4; female TBI, n = 4. One male TBI sample contained two ROIs to adequately capture IBA1^+^ area. One set of ROIs came from a technical replicate slide as shown in (A).

To isolate sex differences associated with TBI rather than baseline differences between males and females, we selected genes that differed between female and male TBI mice (*p* < 0.05, |log□FC| > 1) and were also altered by TBI relative to sham in at least one sex (*p* < 0.05). Using these criteria, we identified 10 genes in the whole somatosensory thalamus and 15 in NeuN-enriched ROIs, with no overlap between the two sets (Fig. 7C–F; Table S2). These sex differences arose primarily from decreased expression after TBI in males or increased expression after TBI in females, although the pattern varied among individual genes (Fig. 7D, F; Fig. S5).

We next asked whether these gene-level differences were associated with broader biological pathways. To identify broad biological pathways across all quantified genes without relying on arbitrary significance thresholds, we performed Gene-set enrichment analysis (*30*). GSEA revealed that female-biased expression was enriched for immune and inflammatory processes, whereas male-biased expression was enriched for synaptic transmission (Fig. 7G, H). These pathway enrichments were more pronounced in NeuN-enriched ROIs. To examine the cellular distribution of these candidate genes, we cross-referenced them with our previously published single-nucleus RNA-sequencing (snRNA-seq) dataset from male mice 3 weeks after CCI (*9*). When detectable, candidate transcripts were most frequently observed in neuronal populations, but expression was also present in oligodendrocytes, oligodendrocyte precursor cells, and astrocytes (Fig. S6). Thus, while NeuN-enriched ROIs enrich for neuronal signals, observed sex-dependent changes may also reflect contributions from non-neuronal cell types.

We then used the snRNA-seq dataset to determine if genes showing a TBI-associated change in males in our GeoMx dataset could be replicated in an independent dataset (*9*). We examined 40 genes showing either sex-dependent gene expression in the GeoMx data or altered expression in males after TBI (Fig. S7). Most candidate genes showed low detection in the snRNA-seq dataset, which may reflect low nuclear transcript abundance. *Clcn4* was upregulated in males but not females after TBI in the GeoMx dataset and was also upregulated in the snRNA-seq dataset. *Clcn4* is an X-linked gene encoding a chloride–proton exchanger associated with intellectual disability and epilepsy in humans (*31*). *Otx2, Slc22a14,* and *Pipox* were three other genes showing conserved, TBI-dependent expression in males across both datasets. These four genes are strong candidates for further study because they demonstrated similar TBI-associated changes in expression across two independent sequencing approaches.

Finally, we examined the spatial distribution of these 40 candidate genes in an independent Visium spatial transcriptomics dataset from male sham and TBI mice (Fig. S8) (*32*). We segmented the thalamus into *Gfap-*high and *Gfap-*low expression to approximate “inflamed” and “uninflamed” thalamic tissue after TBI. Several candidates, including *Rtn4, Uqcrc2,* and *Clcn4*, were expressed at lower levels in the *Gfap-*high than the *Gfap*-low thalamus of TBI mice (log2 fold change –0.37 to –0.49, adjusted *p* < 0.002), but did not differ between corresponding sham regions (adjusted *p* ≥ 0.2), indicating that expression of these candidate genes tracks with regional inflammation (Fig. S8 A–G).

To ask if this regional decrease could be driven by neuronal loss, we estimated neuronal and glial content for each Visium spot using cell-type gene signatures (see Methods). In TBI tissue, the composite neuronal estimate in the *Gfap*-high thalamus was 0.70x that of the *Gfap*-low thalamus, whereas the glial estimate was 2.07x (adjusted *p* < 10^-29^). In contrast, sham regions differed by no more than 1.1x, confirming reduced neuronal density per spot in inflamed tissue near the lesion. Because expression of the canonical neuronal marker *Rbfox3* (NeuN) was low (UMI < 3 per spot) in both sham and TBI animals, we normalized UMI counts to *Snap25*, which was robustly detected across the 208-gene panel (Fig. S8H, I). Normalizing to *Snap25* reversed the regional expression pattern, revealing higher expression of these genes per neuronal transcript in the *Gfap-*high thalamus relative to the *Gfap-*low thalamus (adjusted *p* < 10^-4^). Equivalent reversals were obtained when normalizing to alternative neuronal markers (*Ndrg4, Syt1*, or *Vsnl*). Thus, regional candidate gene expression is TBI-dependent and directly tracks with neuronal transcript density.

Together, these analyses identify sex-dependent molecular responses in surviving thalamic tissue despite broadly comparable neuronal loss. Female-biased expression was associated with immune and inflammatory processes, whereas male-biased expression was associated with synaptic processes. Cross-referencing these findings with independent transcriptomic datasets identified several candidates for further study, with *Clcn4* showing TBI-associated changes across three independent datasets.

## DISCUSSION

Sleep disturbances after TBI differ between females and males, yet most preclinical studies of post-traumatic sleep physiology have focused on males. Here, we found that TBI produced distinct ECoG spectral responses in female and male mice 3 weeks after injury despite broadly comparable cortical lesion size and secondary thalamic neuronal loss. Females exhibited state-dependent changes in delta and mid-gamma activity, whereas males showed broad reductions in ipsilateral spectral power. In contrast, both sexes shared changes in theta organization and sleep spindle properties. Estrous stage further modified a subset of female phenotypes, including mid-gamma activity and spindle peak frequency. Localized sex differences in thalamic C1q expression and neuronal morphology appeared in specific nuclei but did not match the broader structural and histological similarity between sexes. In contrast, exploratory spatial transcriptomics revealed a more global sex-divergent molecular signature in surviving somatosensory thalamic tissue, with female-biased expression linked to immune and inflammatory processes and male-biased expression linked to synaptic transmission. Together, these findings suggest that biological sex shapes the functional and molecular response of surviving neural networks after TBI despite similar tissue and neuronal loss.

### TBI produces distinct electrophysiological phenotypes in females and males

Clinical studies identify female gender as a risk factor for persistent sleep disruption after TBI, but the physiological basis of this association remains unclear (*6, 8*). Our findings indicate that TBI does not produce a single electrophysiological phenotype that differs only in severity between females and males. Instead, specific components of the spectral response to injury differed between sexes and across sleep-wake states. Males showed broad reductions in ipsilateral ECoG power, whereas females showed changes that were more frequency-and state-specific.

The female-specific change in the relationship between NREM and Active Wake delta power is notable because delta power defines NREM sleep and is associated with sleep depth in humans (*33, 34*). Altered delta activity has been associated with seizures following TBI in preclinical models, is affected after TBI in patients, and is observed in patients with clinical insomnia (*10, 35, 36*). Furthermore, delta power shifts during interventions associated with clinical improvement in patients, although whether these changes directly contribute to functional recovery remains unclear.

Females also exhibited state-dependent changes in mid-gamma activity. Gamma oscillations reflect coordinated activity within cortical and hippocampal networks (*37–39*). Altered gamma activity is strongly associated with insomnia and discrepancies between subjective and objective sleep quality in humans outside the context of TBI (*17–19, 40, 41*). While the association between TBI, sleep, and gamma frequencies has not been widely explored, altered gamma frequencies have been reported in patients with TBI (*42*). Together, these findings indicate that sex influences the organization of ECoG activity across sleep-wake states after TBI in frequencies tightly linked to sleep maintenance. Determining whether similar sex-dependent spectral changes occur in patients could clarify whether sex-specific network disruption underlies clinical disparities in post-traumatic sleep symptoms.

Other electrophysiological abnormalities were conserved between sexes. Both females and males showed reduced NREM:Active Wake power ratios in the theta range, reduced spindle magnitude and peak frequency, and impaired interhemispheric spindle coordination. Quantitative EEG studies have similarly identified altered theta activity and reduced spindle magnitude and frequency in patients with TBI (*13, 43, 44*).

Unlike our previous study in male mice, we did not detect a reduction in overall spindle rate after TBI (*9*). Differences in experimental conditions, including daily handling for estrous staging in the present study, could explain this discrepancy, as repeated handling can influence sleep–wake physiology and stress responses (*45, 46*). Nevertheless, the coexistence of shared spindle/theta deficits and sex-divergent spectral responses indicates that biological sex selectively modifies specific components of post-traumatic network activity rather than producing uniformly greater dysfunction in either sex. For instance, while a recent preclinical study reported chronic, male-specific disruptions in sleep–wake macroarchitecture 4–5 months after repetitive blast TBI (*47*), clinical studies associate female sex/gender with worse subjective sleep disturbances (*6, 8*). Our results offer a potential resolution to this discrepancy: female mice exhibit pronounced state-dependent imbalances in ECoG spectral oscillations even in the absence of sex-specific macroarchitectural deficits. This aligns with clinical EEG evidence demonstrating that quantitative alterations in oscillatory sleep rhythms correlate more strongly with subjective sleep quality, daytime sleepiness, and neurological recovery than gross sleep– wake macroarchitecture alone (*17–19, 48–50*).

### Estrous state modifies a subset of post-traumatic electrophysiological phenotypes

The functional consequences of TBI in females also depended in part on estrous state. TBI effects on Active Wake behavior, mid-gamma activity, and spindle peak frequency varied with estrous stage, although the stage-specific pattern depended on the electrophysiological measure. In contrast, the female-specific alteration in the delta NREM:Active Wake relationship did not show a detectable interaction with estrous stage. Thus, ovarian state appears to modify specific post-traumatic electrophysiological phenotypes without providing a general explanation for the overall differences observed between females and males.

Ovarian hormones provide one plausible mechanism underlying sensitivity of TBI effects to the estrous cycle. Estrogen, progesterone, and their metabolites regulate lesion size, molecular markers, and behavioral responses to TBI in preclinical models (*51, 52*). Progesterone administration after diffuse TBI can alter short-and long-term cortical excitability, coinciding with improved sensory and cognitive behavior (*53*). Rhythms affected by the estrous cycle in our data are also reported to vary with the menstrual cycle in humans. In humans and rodents, sleep spindles are sensitive to fluctuations in gonadal hormones (*54–56*), and gamma oscillations can similarly vary with ovarian state (*57, 58*). Fluctuating ovarian hormones can acutely regulate GABAergic and glutamatergic signaling across the estrous cycle, increasing excitability at specific estrous phases (*59–64*). Although this study did not directly measure circulating hormones and estrous stage serves as an indirect proxy for hormone concentrations, a hormonal modulation of the excitation-inhibition balance within local neuronal networks provides a compelling hypothesis for the acute fluctuations observed in post-traumatic oscillations. Other biological factors associated with estrous state or sex chromosomes could also contribute (*65*– *67*). Dissecting how individual hormones impact estrous-dependent sleep-wake rhythms will be essential to establish the mechanisms underlying TBI-disrupted oscillations across diverse hormonal states, including puberty and menopause.

Finally, these findings demonstrate that analyses pooling freely cycling females across estrous stages risk masking hormone-sensitive functional phenotypes. Including both sexes in study cohorts is a critical first step, but resolving hormone-sensitive outcomes requires systematic sampling across individual estrous stages. Emerging technologies enabling real-time, long-term *in vivo* tracking of hormone-responsive neurons (*68*) offer promising avenues to identify the precise circuits and cell types driving estrous-dependent TBI phenotypes, ultimately providing a stronger mechanistic understanding of how gonadal hormones affect TBI outcomes.

### Sex-divergent electrophysiological responses occur despite broadly comparable structural injury

The sex-dependent electrophysiological phenotypes were not accompanied by major differences in structural injury. Cortical lesion was comparable between females and males, as were broad regional measures of C1q and GFAP expression in the cortex and thalamus. Secondary thalamic pathology was strongly nucleus-dependent but generally conserved across sexes. The lateral dorsal thalamus showed substantial neuronal loss after TBI, but the magnitude was comparable between females and males. Thus, neither cortical lesion size nor secondary thalamic neuronal loss paralleled the sex-divergent electrophysiological phenotypes.

However, higher anatomical resolution revealed localized sex differences. C1q expression was greater in male than female TBI mice within the mediodorsal and reticular thalamic nuclei, despite the absence of corresponding sex differences in neuronal loss. C1q participates in complement-dependent synaptic remodeling and contributes to secondary thalamic neurodegeneration and spindle disruption after TBI in male mice (*9*). The reticular thalamus is particularly relevant to spindle generation through its inhibitory interactions with thalamic relay nuclei (*69*). However, the C1q differences did not align with the female-specific spectral phenotypes observed here, arguing against a simple model in which greater complement activation explains sex differences in function.

More broadly, post-traumatic inflammatory responses may differ qualitatively or spatially between sexes rather than being uniformly greater in one sex. Gonadal hormone concentrations are correlated with acute transcriptomic changes in the cortex associated with wound recovery and immune response after TBI (*70*). Sex-dependent microglial states have been reported at the primary injury site 24 hours to one month after TBI in rodent models (*24, 71*), and sex influences microglial function in other neurological disease contexts (*66, 72, 73*). The localized, male-biased C1q differences observed here, together with broadly similar regional C1q and GFAP responses, suggest that sex differences in secondary inflammatory responses may be anatomically and molecularly specific rather than part of a global inflammatory response.

Neuronal morphology showed a similarly localized pattern. Females had smaller neuronal soma size than males specifically in the VPL after TBI, whereas we detected no sex differences in most other thalamic nuclei examined. Although neuronal soma size can covary with intrinsic electrophysiological properties (*74–76*), we did not measure cellular excitability and therefore cannot determine the functional consequence of this morphological difference. Importantly, the VPL phenotype also does not simply explain the electrophysiological findings: spindle abnormalities were largely shared between sexes, whereas the major sex differences occurred in state-dependent spectral activity. Rather, the shared structural vulnerability across thalamocortical relay nuclei—including sex-conserved neuronal loss in LD and soma shrinkage in VPL, VPM, and PO—aligns more closely with the sleep spindle deficits (reduced magnitude, peak frequency, and interhemispheric coordination) observed in both sexes, given the role of dorsal thalamocortical loops in spindle generation and propagation (*77–80*). These observations further argue against a single anatomical lesion or cellular phenotype accounting for the sex-dependent functional response to TBI.

### Surviving thalamic tissue shows sex-dependent molecular responses after TBI

Exploratory spatial transcriptomic analysis further suggested that surviving thalamic tissue responds differently to TBI in females and males. Female-biased expression was associated with immune and inflammatory processes, whereas male-biased expression was associated with synaptic processes. These findings contrast with the histological analyses: regional C1q and GFAP responses were largely comparable between sexes, and the localized C1q differences detected were male-biased. The transcriptomic findings therefore suggest that sex shapes the molecular response within surviving tissue even when overall neuronal survival and inflammation are similar.

These molecular findings should be considered hypothesis-generating. The GeoMx analysis included a limited number of biological replicates, and candidate differential-expression analyses relied on uncorrected *p*-values and effect-size thresholds; no individual genes passed FDR correction. Cross-referencing the GeoMx findings with an independent male-only snRNA-seq dataset supported TBI-associated changes in several candidate genes, including *Clcn4, Otx2, Slc22a14,* and *Pipox*. *Clcn4* was particularly notable because three independent transcriptomic approaches detected a TBI-associated change in males. Analysis of an independent male Visium dataset supported the notion that TBI alters the expression of several candidates, including *Clcn4, Uqcrc2,* and *Rtn4*. However, that dataset contained only one section per condition, and each spot represents mixed cellular populations, precluding between-condition inference and making it challenging to infer cell-type specificity. These datasets therefore provide independent support for candidate molecular responses but do not establish their cellular origin or causal relationship to the electrophysiological phenotypes.

Similarly, the surface ECoG recordings performed here do not localize the precise neural generators of these spectral abnormalities or determine whether subcortical molecular and morphological differences directly drive them. Delta activity can arise from cortical and thalamic sources, whereas cortical gamma activity depends strongly on local inhibitory circuitry (*37–39*), which are themselves modulated by thalamocortical inputs (*81*). Dissecting how subcortical changes contribute to cortical network dysfunction will require targeted, deep-brain electrophysiological recordings combined with cell-type-and sex-specific manipulations.

### Conclusions and implications

Our findings demonstrate that following focal cortical TBI, sex-divergent ECoG spectral dysfunction and molecular responses emerge despite broadly comparable gross lesion size and secondary thalamic neurodegeneration. In contrast, spindle abnormalities were largely shared between sexes, while estrous stage selectively modified specific female electrophysiological phenotypes. These findings underscore the necessity of considering ovarian state and broader sex biology in studies of post-traumatic sleep physiology. Key questions remain about whether these network phenotypes persist, resolve, or evolve during long-term recovery, and which cell types drive state-dependent spectral deficits. Longitudinal studies tracking network function alongside hormone dynamics will be critical to determine whether these sex-dependent physiological alterations predict persistent clinical sleep symptoms, ultimately laying the groundwork for sex-informed therapeutic strategies after TBI.

## MATERIALS AND METHODS

### Experimental design

#### Animals

All animal protocols were approved by the Institutional Animal Care and Use Committee at the University of California, San Francisco and Gladstone Institutes. Adult (6–8-week-old) male and female CD1 mice (Charles River Laboratories, RRID:IMSR CRL:022) were used for all experiments. Animals were group-housed with same-sex littermates and allowed ad libitum access to food and water. Once implanted with devices, mice were individually housed in a facility providing a 12:12 h light/dark cycle (lights on at 07:00).

#### Controlled Cortical Impact Model of TBI

Mice were anesthetized with 3–5% isoflurane and placed in a stereotaxic frame. The center of a 3 mm impactor was aligned to the right somatosensory cortex (-1.0 mm posterior to bregma, +3.0 mm lateral to the midline), and the full impactor was traced with a permanent marker. A dental drill was used to create a craniotomy following the outline of the impactor, and the skull flap was removed. A controlled cortical impact (CCI) was performed with an electromagnetic metal piston (Impact One Stereotaxic Impactor for CCI, Leica Microsystems) using the following parameters: 3 mm flat tip diameter, 20° angle, 0.8 mm depth from the dura, 3.0 m/s velocity, and 100 ms dwell time. For sham surgeries, mice received skull thinning with a dental drill over the same 3 mm diameter region but received no impact and maintained an intact skull.

#### Surgical Implantation of Wireless Telemetry Devices for Chronic ECoG Recordings

Three stainless steel screws were soldered using lead-free solder (Kester; 24-7068-6417) and solder flux (Harris; 1UYK3) to leads on wireless telemetry devices from Data Sciences International (HDX-02). References for both leads were soldered to the same stainless-steel screw as a single reference electrode. Completed devices were left in 1% Tergazyme (Electron Microscopy Sciences; 60520) overnight prior to surgery and briefly rinsed in deionized water immediately prior to implantation.

Mice were implanted with the above devices two weeks following the TBI surgery. Briefly, mice were anesthetized with 3–5% isoflurane and placed in a stereotaxic frame. A dental drill was used to drill small holes for ECoG screws in the ipsilateral (+0.5 mm anterior to bregma, +3.0 mm lateral to midline) and contralateral (+0.5 mm anterior to bregma,-3.0 mm lateral to midline) somatosensory cortex. Electrodes were screwed in the minimum number of turns to stabilize the screw. The ground/reference electrode was placed slightly more posterior and right of lambda in the cerebellum. The battery/transmitter device was placed under the skin in the mouse’s abdomen. The surface of the skull, including the TBI/sham region, was covered with a small layer of dental cement to secure and insulate the screws, and the skin was sutured. Mice were singly housed and allowed to recover for one week before recording.

One week after device implantation (three weeks after TBI/sham surgery), each home cage was placed over a wireless receiver, and ECoG signals for up to eight mice were simultaneously recorded for a consecutive 4-6 days using the Ponemah Software (DSI v. 6.5.1) and sampled at 500 Hz.

#### Estrous Staging

Female mice had daily vaginal lavages performed between zeitgeber time (ZT) 2 and ZT 4. For vaginal lavages, 25 μL of room-temperature 1× PBS was washed into the vaginal canal using a pipette. Samples were deposited onto a slide and allowed to dry completely before Giemsa staining. Males underwent the same handling to control for handling stress.

#### Giemsa Staining

A stock Giemsa stain was prepared as described previously (*82*). Briefly, a stock Giemsa stain was prepared at least one week prior to staining. An 18.5% solution of Giemsa powder was prepared in glycerin, heated to 60°C, and cooled. This solution was then diluted 9:14 in 100% methanol to create a stock solution.

Slides with dried samples were briefly dipped in 100% methanol for fixation. Stock Giemsa solution was diluted 1:30 in PBS and shaken vigorously. Slides were incubated in Giemsa solution for ∼1.5 hours, dipped in distilled water to rinse, and cover-slipped. Using light microscopy, estrous stage was determined by the relative abundance of leukocytes, nucleated epithelial cells, and cornified epithelial cells. If an estrous stage was challenging to call, high-confidence estrous stages prior and after the date were used to inform estrous stage assignment. If no high-confidence estrous stages were available to make the decision, or the assignment remained unclear, we excluded the data point. To keep our results consistent with ZT alignment, we assigned morning estrous staging results to the day they occurred. However, we acknowledge that behavioral estrous typically occurs in the middle of the dark phase prior to the estrous phase captured in the morning (*83*), and therefore do not attempt to make any conclusions on the precise hormones involved in these processes.

### Analysis of ECoG Recordings

#### Sleep Staging

Data were visually inspected using Spike2 (CED, v8.21+), and channels with high noise contamination were excluded from downstream analyses. Data from our DSI system (4–6 day continuous recordings) were chunked into 4-hour blocks for processing with a 15-second overlap across chunks. Major artifacts were defined as events > +0.75 mV and < –1.0 mV ± a minimum of 2 seconds surrounding the event was excluded from the data. Precise boundaries for the artifacts were chosen to minimize artificial jumps in the data. Data were processed using multitaper spectrogram analysis functions developed by the Prerau Lab using mean-subtraction detrending, a window of 30 seconds, step size of 1 second, 29 tapers, and a time-bandwidth product of 15 (*84*). Delta (2–4 Hz), Theta (6–8 Hz), and Mid-Gamma (60–80 Hz) were summed across the multitaper results to create a single summary value for sleep scoring with a time resolution of 1 second. We chose a narrower delta band because low-frequency, high-amplitude events were difficult to remove from the data. A narrower theta band was also used here, as this definition performed better at consistently segmenting out REM/Quiet Wake than a 6–9 Hz theta definition. As our devices did not have EMG, we estimated sleep states using two sequential 2-state HMM models to first segment wake from sleep and then to segment sleep into NREM and short arousals/REM. The first HMM model contained features initialized on z-scored delta and mid-gamma power. The creation/fit of the model was run 50 times and averaged across iterations to generate a probability of being in “sleep” or “wake” for a single 1-second window. Iterations where state means were greater than 0.75 standard deviations apart and delta/mid-gamma means had the same sign, or iterations where state means were < 0.5 standard deviations apart, were determined as non-discriminatory and removed from state estimations. After scoring, bouts of sleep or wake less than two minutes were flipped to align with the previous stage to ensure enough quiet wake/REM was present in sleep to segment out. Next, epochs labeled as sleep were run through another 2-state HMM model initialized on z-scored delta and z-scored theta:delta ratio to identify putative REM bouts, seeded with a transition matrix biased toward state stability, and microarousals using the same HMM methodology described above. These putative REM/microarousal bouts were relabeled to “Wake/REM” and bouts 1 minute or less were switched to “NREM”. As we did not have an EMG, our activity counts were primarily sensitive to ambulatory movement, and our TBI mice exhibited altered theta; we found it challenging to confidently separate quiet wake bouts from REM in TBI mice and grouped these stages together. All data were manually validated using a custom GUI.

Following sleep scoring, the total amount of movement for a single stage was calculated using the DSI system’s activity counts metric, which uses a threshold-based change in device signal strength to register a single “Activity” count. To classify data as “Quiet” or “Active” Wake, the 95th percentile across Activity Rates during NREM bouts was used as a threshold and manually checked against the density distribution of Activity Rates for NREM and Wake/REM on a per-animal basis. The 90th percentile of NREM activity rates was chosen as a threshold for one animal with higher overlap between the NREM and Wake/REM distributions of Activity Rates.

For statistical analysis on per-day data, we used linear mixed-effects models on the per-animal means for bout duration, number of transitions, and percent time with TBI status, sex, and their interaction as fixed effects and a random intercept for animal ID. Model fits were assessed using residual diagnostic plots, and significance of fixed effects was assessed using Type III ANOVA with Satterthwaite’s estimation (lmer, lmertest). Pairwise comparisons were performed using estimated marginal means (emmeans) to test for a TBI effect or sex effect across all groups. Sex differences were only analyzed post hoc if sex was a significant main effect or covariate. *P*-values were adjusted using the Holm-Bonferroni procedure for all post hoc tests conducted for each outcome measure, as the number of comparisons was small and fixed.

To assess time-of-day effects, we calculated per-hour summaries of bout number, mean bout duration, and proportion of time for each recorded day and used linear mixed-effects models. TBI status, sex, and ZT hour were set as fixed effects, including interactions, and animal ID was set as a random effect. Model fit and significance of individual variables were assessed as described above.

To assess estrous cycle effects, we fit new linear mixed-effects models on the per-day means for bout duration, number of transitions, and percent time with TBI status and estrous stage as fixed effects, including interactions, and animal ID as a random intercept. Multiple comparisons were corrected for using the false discovery rate (FDR) method for all post hoc comparisons, as this was a larger comparison family.

#### Bandpass calculations for power spectral analyses

To calculate power across bandpass frequencies, we again performed multitaper spectral analysis as described above, instead using a 5-second window, 1-second steps, 4 tapers, and a time-half bandwidth of 2.5. For ipsi/contra analysis, the ipsi/contra ratio was calculated by taking each individual vigilance bout and averaging the power for each frequency bin per electrode to get one ipsi/contra ratio per vigilance bout. These values were then averaged across all bouts, and the log-ratio between ipsilateral and contralateral side was calculated within NREM, Active Wake, or Quiet Wake & REM. To calculate the NREM-to-Active Wake ratio, power in each frequency bin was first averaged within a bout, then across all bouts belonging to NREM and Active Wake, and the log-ratio was calculated between NREM and Active Wake. We excluded mice with an ipsilateral-to-contralateral ratio more than 2 standard deviations above the group mean in 6 of 7 frequency bands post-analysis to remove systematic outliers, and we manually checked data quality in the ipsilateral and contralateral EEG. This removed two male TBI animals from the ipsilateral-to-contralateral normalization analysis, as manual examination of their EEG showed lower-quality signal from the contralateral channel. As individual rhythms were still present, we included data from these electrodes in analyses that performed within-electrode normalization.

For the ipsilateral-to-contralateral ratio, we fit a single linear mixed-effects model per frequency band instead of one overall model based on the quantile-quantile (Q-Q) plots. Each frequency-band model included TBI status, sex, and vigilance state as fixed effects, including all interactions, and a random intercept for animal ID to account for repeated measurements within each mouse. Model fit was assessed by inspecting residuals, Q-Q, and random-intercept plots. Statistical significance of fixed effects was assessed using Type III ANOVA with Satterthwaite’s approximation for degrees of freedom (lmer, lmertest). Pairwise comparisons of condition within sex and behavioral state were conducted using estimated marginal means (emmeans). Post-hoc sex comparisons were only calculated if sex was a significant main or interaction effect in the linear model. *P*-values were corrected for multiple comparisons across sex, TBI status, and vigilance state using the FDR procedure for all post hoc comparisons.

For NREM to Active Wake Ratios, we used one linear mixed-effects model with Condition, Sex, Band, and Side as fixed effects, included all interactions, and set a random intercept for each animal ID to account for repeated measurements within a mouse. Model fit, post-hoc comparisons, and multiple-comparisons correction were assessed as described above, except multiple comparisons were corrected within each band for TBI and sex comparisons across both hemispheres (ipsilateral and contralateral) rather than vigilance states.

For the estrous cycle, per-animal averages were calculated within each estrous stage for female animals and modeled using the same strategy as above, with TBI status, estrous stage, and hemisphere included as fixed effects in the ipsilateral to contralateral ratio and TBI status, estrous stage, and frequency band as fixed effects in the NREM to Active Wake ratio. Animal ID was included as a random effect in both models, and the same methods for evaluating significance of fixed effects, pairwise comparisons, and model fit were used in this section.

#### Sleep Spindle Detection

Discrete sleep spindles were detected using a Morlet Wavelet detection method as previously described (*9*). Briefly, a Morlet wavelet with a central frequency of 13 Hz and 12 oscillations was convolved with all segments labeled as NREM in the raw ECoG. We also tested a Morlet wavelet with a central frequency of 13 Hz and 7 oscillations and identified similar results. Spindles were defined as events crossing an upper threshold (mean + 1.5*SD) and spindle edges were identified as the nearest crossings of a lower threshold (mean + 0.5*SD). Spindles were limited to events between 0.5–3 seconds long based on standard definitions (*85*). A random subset of spindles was validated by eye for each mouse. Spindle rates were calculated for each bout of NREM and normalized by the duration. NREM bouts with no spindles detected were excluded from spindle rate analyses as the statistical fit was better on spindle rates for NREM bouts with just spindles. We also analyzed the proportion of NREM bouts without spindles and ran a spindle-rate analysis that included NREM bouts with no spindles in the overall spindle rate, and found no TBI effect in either outcome.

For each spindle, the magnitude, peak frequency, and duration were calculated. To calculate magnitude, we first applied a 3rd-order Butterworth filter to isolate the 10–15 Hz signal. We then averaged the absolute value of the band-passed signal. The peak frequency of the spindle was determined by performing a fast Fourier transform on the spindle and selecting the largest peak between 10–15 Hz as previously described (*86*).

To identify overlapping spindles, we used the foverlaps function in R and calculated the percent time a spindle overlapped with a co-occurring spindle on the opposite cortex by dividing the overlap duration by the spindle duration. Spindles that shared at least 25% of their total duration with a spindle in the opposite hemisphere were considered paired. The normalized value per mouse was calculated by taking the number of overlapping spindles divided by the number of unique spindles across the ipsilateral and contralateral cortices.

For statistical analyses, we used a linear mixed-effects model on per-animal averages for each spindle measure (magnitude, peak frequency, duration, rate, spindle overlap), with TBI status, sex, and hemisphere as fixed effects (including all interactions) and a random intercept for each animal. For spindle overlap, condition, sex, and their interaction were included as fixed effects. For spindle overlap split by direction (ipsi overlapping with contra; contra overlapping with ipsi), direction was also included as a term. Models were fit using lmer from the lme4 package, and fit was evaluated using residual diagnostic plots. Type III ANOVAs with Satterthwaite approximation were used to assess the significance of fixed effects. Pairwise comparisons between condition and hemisphere were performed using estimated marginal means with Holm-Bonferroni adjustment of *p*-values within each hemisphere. Post-hoc investigations of sex were not included as sex was not a significant main or interaction effect in any model.

For the estrous cycle, per-animal averages were calculated within each estrous stage for female animals and modeled using the same strategy as above with TBI status, estrous stage, and hemisphere included as fixed effects and animal ID as a random effect. FDR was used to correct *p*-values across both hemispheres for measures showing a significant estrous cycle effect.

### Histology

#### Tissue Processing

Mice were anesthetized with a lethal dose of Fatal-Plus and perfused with chilled 1x PBS then 4% PFA in 1x PBS. Brains were left in 4% PFA overnight, then transferred to 30% sucrose for at least 3 days. We took topographical images of the lesion at 2.5x using the Keyence Profilometer (VK-X1000). Two male sham and two male TBI animals were discarded due to poor fixation. We then cut serial brain sections at 40 μm thick on a Leica SM2000R sliding microtome. For all histology, we used 4–5 sections per animal spanning the thalamus spaced at ∼ 280 μm apart, including 1–2 extra sections outside of our target area for angle adjustment and alignment to the Allen Brain Atlas.

#### Lesion Analysis

In the Keyence MultiFile Analyzer software (v. 2.2.0.93), we calibrated the reference plane using manually set points in a circle approximately 3 mm in diameter (the size of our impactor). If tissue damage was visible on the brain surface, we manually placed points in areas surrounding the injury that did not show obvious edema or tissue loss. Quantification of the lesion was restricted to either a ∼3 mm circle (if tissue damage was not visible) or manually traced (if tissue damage was visible). The tissue displacement above and below the reference plane (edema and cavity) were quantified separately, and then the absolute values of both numbers were added to get total tissue displacement. The average of this value was then calculated separately for male and female sham animals. Each animal’s tissue displacement value was then divided by the sex-matched sham average value. Statistics were calculated using the two-sided Wilcoxon rank-sum test with Holm-Bonferroni correction for multiple comparisons.

#### Immunofluorescence staining

We washed sections 3x with PBS and permeabilized for 10 minutes in PBS with fresh 0.5% Triton. Sections were washed 3x for 5 minutes with 0.05% Triton in 1X PBS (PBST). Sections were then blocked for 1 hour with 10% normal goat serum diluted in PBST and then incubated overnight at 4°C with antibodies directed against C1q (1:500; Abcam, AB182451)/Gfap (1:1000; Invitrogen 13-0300). After washing, sections were incubated with the following Alexa-Fluor-conjugated secondary antibodies (C1q/Gfap: AF594, Goat anti-rabbit, A11012, Lot#: 1844440/AF488, Goat anti-rat, A11006, Lot#: 1887148) for 2 hours. Sections were washed 3x in PBST, then 1x in PBS before being mounted with VectaShield and coverslipped. Slides were imaged at 2x magnification on a Keyence BZ-X800 microscope.

#### 3,3’-Diaminobenzidine staining

We washed sections 3x for 5 minutes in PBS + 0.1% Tween 20 (PBSTw) before incubating them in 1% H2O2 + 30% ethanol+ PBS for 20 minutes. After washing sections 3x for 5 minutes in PBSTw, we blocked them in 20% normal goat serum (Jackson Immunoresearch Laboratories, Lot #170983) + Mouse anti-mouse (2 drops per 2.5 mL, MKB2213-1) blocking reagent + PBSTw for 1 hour. We then incubated sections with NeuN (1:500, Sigma-Aldrich, MAB377) in 3% normal goat serum + PBSTw. Sections were washed in PBSTw 3x for 5 minutes, incubated with Mouse Mach2 Polymer (Biocare Medical, MHRP520L) for 1 hour, washed with PBSTw 3x for 5 minutes, and then developed with a DAB kit (Vector Lab, SK-4100) for ∼1 minute and 45 seconds. Sections were washed in PBS 1x for 5 minutes and mounted on SuperFrost Plus slides. To dehydrate sections, slides were placed in 70% Ethanol for 1 minute, 95% Ethanol 2x for 1 minute, 100% Ethanol 2x for 2 minutes, and then placed in Xylene 2x for 2 minutes. Sections were then mounted in Shandon Mount and coverslipped. Slides were imaged using brightfield imaging at 20x magnification with a Leica Versa Slide Scanner.

#### Alignment to Allen Brain Atlas

For immunofluorescent stains, only the DAPI channel was used for alignment, and all images were calibrated to the BZ-X800 manufacturer’s specifications using a custom FIJI macro. All DAPI and chromogenic NeuN images for one mouse were imported into the Aligning Big Brains & Atlases (ABBA v.0.80) using BioFormats. Chromogenic NeuN^+^ sections were used to adjust the angle of the reference atlas before aligning immunofluorescent stains to the 3D Mouse Allen Brain Atlas. Adjacent sections from different stains were placed approximately 40 μm apart. Each section was registered to the reference atlas using BigWarp, and all regions of interest (ROIs) were exported to an ImageJ-compatible format. Sections that had poor-quality staining or had significant obstruction of the thalamus were excluded from downstream analyses.

#### Fluorescence Quantification

For each section, we used custom FIJI macros to manually outline and exclude torn, folded, or blurry regions from downstream analyses in the DAPI channel, split ROIs into ipsilateral and contralateral sides, and measure integrated density, raw integrated density, and area for a selected set of thalamic nuclei in fluorescent channels corresponding to C1q and Gfap. Fluorescent density was quantified for both the ipsilateral and contralateral sides by dividing the raw integrated density by the area of the ROI. A fluorescence density ratio was created by dividing the fluorescent density on the ipsilateral side by the same region on the contralateral side. Fluorescence density ratios were averaged across all included sections to get a single average ratio per mouse. Any sections that did not have the region present on both the ipsilateral side and contralateral side were excluded from analyses. Percent area was determined by semi-manually thresholding the image to match the visible signal by an experimenter blind to condition and intersecting the thresholded area with each ROI. The percent area was calculated as the total area of the thresholded stain within an ROI divided by the ROI area. Two-sided Wilcoxon-rank sum tests with FDR-correction were used for statistics as there were multiple outliers when fitting with linear mixed-effects models, resulting in poor model fit. Correction for multiple comparisons was performed individually for each protein (GFAP or C1q) analyzed across all brain regions quantified.

#### Detection and Quantification of NeuN^+^ Cells

For each thalamic region of interest, we generated a cropped image and mask corresponding to the ROI. Using CellProfiler (4.2.8), we estimated the background signal using a Gaussian filter with sigma = 15 and subtracted it from the same image smoothed with a Gaussian filter with sigma = 5. From this image, we used the Global Otsu Three-class method of identifying neurons in the IdentifyPrimaryObjects module (typical diameter between 17–2000 pixels, including objects touching the border of the image, assigning the middle intensity class to the foreground, threshold smoothing scale of 1.3488, declumping filter of 15, minimum distance of 15). Putative neuron centers were excluded if the neurons were too thin (minor axis length under 25 pixels, eccentricity greater than 0.9, and size less than 500 pixels). These parameters were chosen by using the EditObjectsManually module to manually outline neurons on a small subset of images and comparing automated and manual detection of neurons. Parameters were chosen based on visual cues and PCA-based methods to identify features with major discriminatory power. We then calculated cell size in um using the microscope’s pixel-to-um scale and applied a final cell-size filter to limit cells to 5–200 μm^2,^ removing at most 2% of cells from each sex-condition pairing. Neuronal densities (cells per mm^2^) were calculated by dividing the number of NeuN^+^ cells by the tissue area within each brain region per section and averaging across all sections.

Linear mixed-effects modeling was used on neuronal density and median cell size. We fit an LMM with TBI status, sex, and brain region as fixed effects, including all interactions, and animal ID as a random effect to account for repeated sampling. Models were fit using lmer, assessed for fit using residual diagnostic plots, and evaluated for significance of fixed effects using Type III ANOVA with Satterthwaite’s approximation and estimated marginal means. P-values were corrected for multiple comparisons using FDR correction across all regions analyzed. We first tested for TBI effects and, if significant, we performed a secondary test to examine sex differences within the TBI group, correcting for multiple comparisons across the regions tested.

### Spatial Transcriptomics

#### Sample Preparation

Mice were given TBI or sham surgeries as described above and perfused with RNase-free 10x PBS diluted in RNase-free, DEPC-treated water (Water: ChemCruz 7732-18-5, 10x PBS: Corning 46013-CM), then 4% PFA, and transferred to 30% sucrose for 24 hours. Brains were then embedded in Cryogel and placed at-80°C. Mice were randomly assigned to 4 groups so that each slide had 1 male TBI, 1 male sham, 1 female TBI, and 1 female sham section located within the capture region. For each group, we used a Leica cryostat (Leica Microsystems) to collect sequential coronal brain sections spanning the thalamus, 5 micrometers thick, one mouse at a time, taking care to ensure each section was at approximately the same bregma. One group had two sections outside the capture region, so a technical replicate was included in place of that group, leaving 3 independent groups and one technical replicate.

#### Digital spatial profiling of mouse brain tissue

Cryogel-embedded mouse brain samples were profiled using GeoMx^®^ DSP (*87*). Thin (5 µm) tissue sections were prepared according to the manufacturer’s recommendations for the Semi-Automated RNA Slide Preparation Protocol (FFPE) (manual no. MAN-10151-06). Four slides containing mouse coronal brain sections were baked for 3 hours at 65°C before deparaffinization, rehydration, heat-induced epitope retrieval, and enzymatic digestion. Tissues were then incubated with 10% neutral buffered formalin (NBF) for 5 minutes and 5 minutes with NBF Stop buffer. All steps following baking were carried out on a Leica BOND-RX. Slides were then removed from the Leica BOND-RX, and *in situ* hybridization with GeoMx^®^ Mouse Whole Transcriptome Atlas probes was carried out overnight in a humidified hybridization chamber kept at 37°C. Following rounds of stringent washing with a 1:1 volumetric mixture of 4x Saline Sodium Citrate buffer (4x SSC) and 100% formamide to remove off-target probes, the tissue was blocked with Buffer W blocking solution (Bruker Spatial Biology), then incubated with a cocktail of anti-NeuN-AF488, anti-GFAP-AF594, and anti-IBA1-AF647 antibodies for 60 minutes at room temperature. The staining cocktail also included the nuclear marker Syto83.

Tissue morphology was visualized using the fluorescent antibodies and Syto83 on the GeoMx^®^ DSP instrument. ROIs were selected from the samples using the polygon or geometric tool. ROIs were either collected as full ROIs or segmented to capture NeuN^+^, IBA1^+^, and NeuN^-^/IBA1^-^ areas. Two male TBI animals had two ROIs in the thalamus collected in order to adequately capture the IBA1 signal. One of these extra ROIs included both IBA1^+^ and NeuN^+^ ROIs, and the other was composed only of IBA1^+^ regions and IBA1^-^ regions. For two female sham animals, the contralateral side of the brain was used due to folds on the ipsilateral thalamus. UV light was then utilized to release and collect oligonucleotides from each ROI. Areas of UV light irradiation within an ROI are referred to as areas of illumination (AOI). Detailed methods for library construction can be found in the Bruker Spatial Biology GeoMx® DSP NGS Readout User Manual (manual no. MAN-10153-06). Briefly, during PCR, Illumina i5 and i7 dual-indexing primers were added to each photocleaved oligonucleotide, allowing for unique indexing of each AOI. Library concentration was measured using a Qubit fluorometer (Thermo Fisher Scientific), and quality was assessed using a Bioanalyzer (Agilent). The Illumina NovaSeq 6000 was used for sequencing, and the resulting FASTQ files were then processed by the NanoString DND pipeline to generate count data for each target probe in every AOI.

#### Quality control and preprocessing of GeoMx transcript data

The GeoMx DSP Analysis Suite was utilized for conducting both quality control (QC) and data exploration. First, each AOI was QC-checked to ensure contamination was avoided during PCR and library preparation and that sequencing was sufficient. Prior to normalization, any genes not detected above background, as determined by the expression of a set of negative probes, in at least one AOI were filtered out from the dataset. The remaining genes were normalized using third quantile (Q3) normalization. The QC-checked, Q3-normalized dataset was used for downstream analysis.

We used linear mixed-effects modeling to examine the interaction between sex and TBI status on probe counts across ROIs, with animal ID as a random intercept to account for nested data. Models were fit using the lmer function from the lme4 package in R. Statistical significance of fixed effects was determined using a Type III ANOVA with Satterthwaite’s approximation. To evaluate simple effects, we used estimated marginal means to test female TBI v. male TBI, female sham v. male sham, female sham v. female TBI, and male sham v. male TBI.

To identify candidate genes for exploratory profiling, differential expression was evaluated using uncorrected p-values (*p* < 0.05) and effect-size thresholds (|log_2_-fold change| > 1), as no individual genes survived genome-wide FDR correction. We identified candidate genes as sex-divergent if they exhibited *p* < 0.05 and |log_2_-fold change| > 1 between male and female TBI animals, and were also significantly altered by TBI relative to sham in at least one sex (sham v. TBI *p* < 0.05 for either males, females, or both). This filtering was performed for the whole ROI (all immunostaining groups combined) and for the NeuN-enriched ROI (NeuN^+^ immunostaining). Next, we performed Gene Set Enrichment Analysis (GSEA) using gseGO from clusterProfiler v. 4.18.4 (*30*). Briefly, genes were ranked based on the sign of the log_2_-fold change and raw *p*-value and then submitted for GSEA using the Biological Process (BP) subset of the Gene Ontology Database. Enriched GO terms were corrected for multiple comparisons using Benjamini-Hochberg correction.

#### Validation in Independent single-nucleus RNA-seq Dataset

We used the R object from Holden et al. (*9*) (GSE179541) as a second-method validation of our GeoMx results. Genes were included in this analysis if: 1) they were significantly different between male and female TBI animals and were significantly different between sham and TBI male mice, or 2) they were significantly different between sham and TBI males and had an effect size corresponding to |log-fold change| > 1. We used significance in males as a primary filter as the snRNA-seq dataset contained only male data. For our NeuN-enriched region, we restricted analyses to glutamatergic and GABAergic neurons and compared expression between sham and TBI animals using the two-sided Wilcoxon rank-sum test in Seurat, with *p*-values adjusted for multiple comparisons across all genes using Benjamini-Hochberg correction.

#### Spatial Analysis of Independent Visium Dataset

We used the Visium spatial transcriptomics dataset from Cho et al. (*32*) as an independent, region-resolved test of the candidate genes nominated by our GeoMx analysis (Fig. 7C–F)(*32*). The analyses used one sham and one TBI hemisection from male mice on a C57BL/6J background profiled six weeks after surgery (2,321 and 2,214 spots in tissue, respectively). We reused the ROIs defined in that study rather than drawing new ones: the thalamic ROIs correspond to the two clusters identified there by unbiased k-means clustering of all spots into ten clusters, which separated into *Gfap*-high (proximal) and *Gfap*-low (distal) thalamic territories, and the cortical ROIs correspond to cortex proximal and distal to the lesion. Spot barcodes were matched to these ROI annotations and to array coordinates, which were converted to physical distances using the Visium v1 geometry (50 μm between array columns and 86.6 μm between rows). Spots were registered to the hematoxylin-and-eosin images, and the array indices were confirmed to match the published coordinate file, accounting for the expected one-based offset.

Expression was analyzed as raw UMI counts per spot, following the source study. For each gene, we compared the proximal and distal ROIs of each structure within each section, using all spots in the ROIs, including those with a count of zero. A comparison was tested when the two ROIs together contained at least 3 spots with a positive count, which gave 83 testable comparisons of the 160 possible (40 genes x 2 structures x 2 sections). Genes were additionally grouped into four detection tiers based on their detection rate across ROI spots and testable ROI pairs: *Well powered*: n = 8 genes, detected in 23–99% of ROI spots, all four ROI pairs testable; *Testable*: n = 8 genes, detected in 3–10% of ROI spots, all four pairs testable; *Sparse detection*: n = 16 genes, detected in 0–7% of ROI spots, 0–3 pairs testable; and *Not detected*: n = 8 genes, 0 positive spots in either section.

Group differences were assessed with two-sided Wilcoxon rank-sum tests, and p-values were adjusted across all 83 testable comparisons using the Benjamini-Hochberg procedure; an adjusted *p* < 0.05 was considered significant.

Because the proximal ROIs lose neurons and gain glia after injury, a change in counts per spot can reflect the number of neurons sampled rather than expression per neuron. Each spot contains several cell types, so we first asked which genes rise and fall together across spots of the sham section: non-negative matrix factorization was used to express the spot-by-gene count matrix as eight gene groups, each defined by a weight per gene and an amount per spot. Six of the eight groups were dominated by neuronal genes and corresponded to the neuronal populations present in the section (cortical, hippocampal, thalamic, striatal, *Pvalb*-rich and *Gap43*-rich); one was dominated by oligodendrocyte genes and one by astrocytic genes; eight was chosen as the smallest number that resolved these neuronal territories while keeping the oligodendrocyte and astrocytic groups distinct. A gene was assigned to a group when its weight there exceeded 1.5× its weight in the others. A gene entered the neuronal set only if it was highly weighted in all six neuronal groups, so the set contains genes that neurons express everywhere in the section rather than markers of one region; the glial and oligodendrocyte sets each came from their one corresponding group. Scoring every spot with the resulting neuronal and glial gene sets, the neuronal score in the proximal thalamic ROI of the TBI section was 0.70× that of the distal ROI, while the glial score was 2.07× (adjusted *p* = 9.8 × 10□³□ and 7.1 × 10□□□), against 0.95× and 1.10× for the same comparison in the sham section, confirming that neuronal content differs between the regions being compared.

Candidate normalizing genes had to be measurable in essentially every spot of both sections, defined as detection in ≥ 98% of ROI spots with a mean of ≥ 8 UMI per spot; 208 of 20,476 genes satisfied this criterion, whereas *Rbfox3* (NeuN) did not (detected in 66% of ROI spots at 1.2 UMI per spot). These 208 genes were ranked in the sham section by how specific they were to neurons: (1) Expression in gray matter relative to white matter (defined per spot using the oligodendrocyte gene set); (2) Correlation with the neuronal vs. glial score; and (3) The degree to which they eliminated apparent neuron-loss signal from known pan-neuronal genes.

*Snap25* ranked second of 208 for neuron specificity, behind *Vsnl1*, and removed 85% of the neuron-loss signal, compared with 28% when counts were normalized to the total per spot.

Four pan-neuronal genes met all criteria: *Snap25, Ndrg4, Syt1*, and *Vsnl1*. *Snap25* was used because it carries the most counts of the four (40.6 UMI per spot, against 10.8–32.2) and therefore gave the narrowest bootstrap 95% confidence intervals across the reported effects (mean width 0.77, vs. 0.84 for *Ndrg4*, 1.10 for *Syt1* and 1.11 for *Vsnl1*).

Candidate-gene raw counts were divided by *Snap25* counts in the same spot, excluding the few spots with no *Snap25* count (2 of 46 spots in one ROI, none elsewhere), and proximal and distal ROIs were compared as above, with *p*-values adjusted across the 24 normalized comparisons. Normalizing instead to *Ndrg4, Syt1*, or *Vsnl1* gave the same direction of effect for every reported comparison (log□fold change +0.52 to +1.04 in the injured thalamus); *Ndrg4* is neuron-restricted without being a synaptic transcript, so this agreement does not depend on the normalizing gene belonging to the synaptic machinery, as the other three do.

Analyses were performed in Python 3.11 using NumPy, SciPy, statsmodels, and scikit-learn. Because this dataset contains a single section per condition, all comparisons were made within sections, and no statistical comparisons were made between sham and TBI tissue.

### AI Usage Statement

Claude Opus 4.7+ was used to reorganize, comment, and draft code. Gemini Pro 3.1, Gemini Thinking 3.5, Claude Sonnet 4+, and Claude Opus 4.7+ were also used to troubleshoot code. All code was tested and validated by the first author to ensure accuracy and reproducibility. During drafting of the text, Claude Sonnet 4+ was used to improve grammar and clarity. AI-generated suggestions were reviewed and revised by the first and last author.

## Acknowledgments

We thank Loren Frank and Eric Denovellis for advice on ECoG analysis, Jonathan Meng for advice on Cell Profiler Analyses, Bret Holt for help with animal husbandry, and Reuben Thomas from Gladstone’s Bioinformatics Core for helpful advice on statistics. We also thank Mercedes Paredes and Daniele Canzio for use of their cryostat and the Nanostring/Bruker TAP teams (Sarah Church, Ben Sutton, Kara Gorman, and Michael McKenna) for spatial transcriptomics and a detailed methods section. We thank Francoise Chanut for help with scientific editing and Tami Tolpa for help with graphical design. We thank Shreyaa Gunasekar, Parinita Saluja, Shravan Kannan, Emily Axelrod, and Zoey Ruderman for careful review and feedback on the manuscript.

## Author contributions: CRediT

Conceptualization: A.M.M., J.T.P.

Methodology: A.M.M., Y.V., D.W., F.S.C., J.T.P.

Data Curation: A.M.M., Y.V., F.S.C., J.T.P.

Investigation/Formal Analysis: A.M.M., Y.V., F.S.C., J.T.P.

Validation: A.M.M., A.C., J.T.P.

Visualization: A.M.M., Y.V., J.T.P

Supervision: A.M.M., J.T.P.

Writing—original draft: A.M.M., Y.V., J.T.P.

Writing—review & editing: A.M.M., Y.V., J.T.P.

Funding acquisition: A.M.M., J.T.P

Resources: J.T.P.

## Data, code, and materials availability

ECoG Data will be posted on the DANDI database upon publication, and code used to analyze data will be archived on Zenodo upon publication and available through a GitHub repository. Raw GeoMx Matrices will be deposited into GEO upon publication. All other data needed to evaluate and reproduce the results in the paper are present in the paper and/or the Supplementary Materials. This study did not generate any new materials.

## Supplementary Materials

**Fig. S1.**
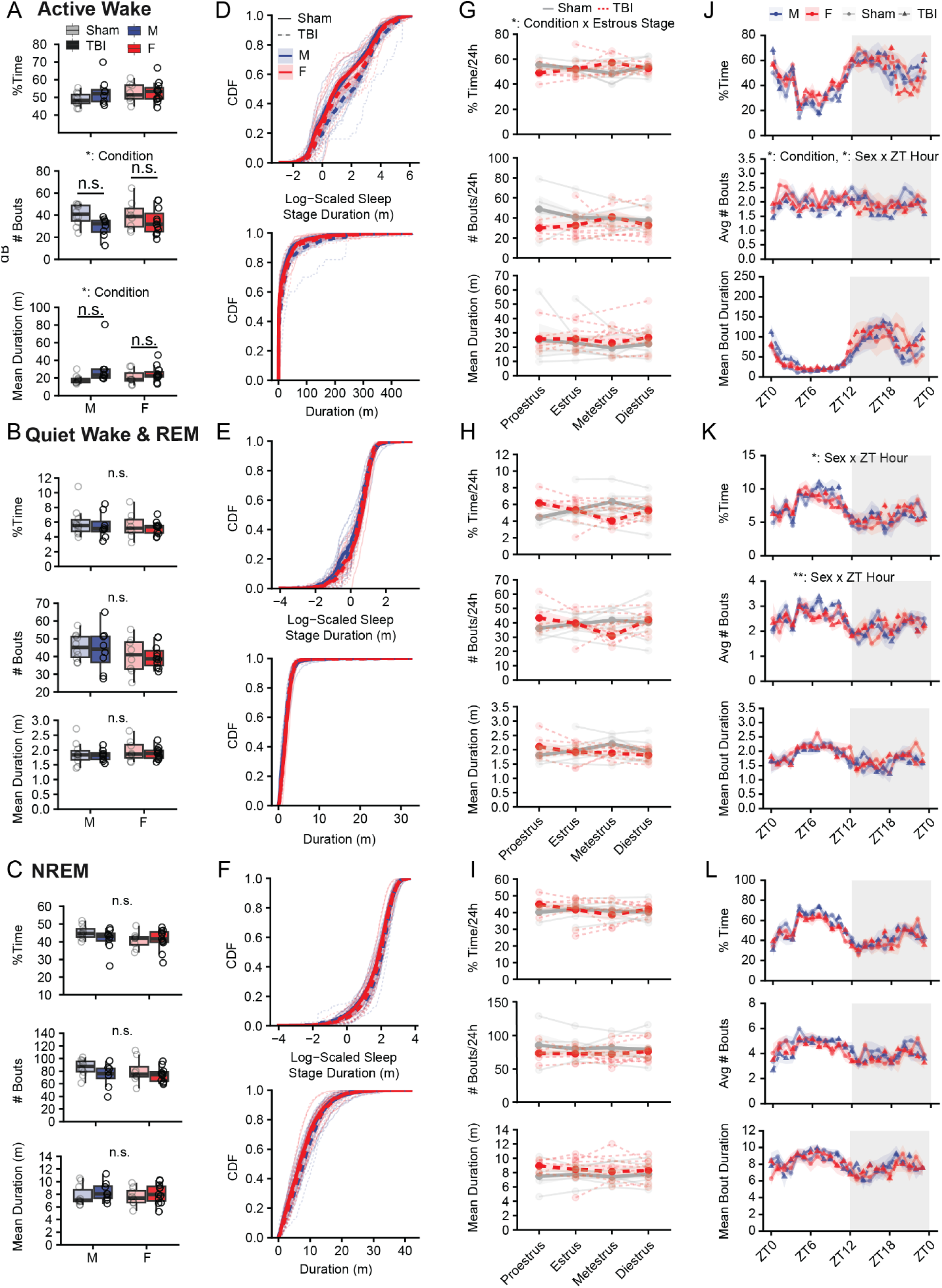
TBI alters Active Wake temporal organization in both sexes, with estrous-dependent effects in females. (A) Percentage of time, number of bouts, and mean bout duration for Active Wake over a 24-hour period. Each point represents the mean value for one animal across 4 recorded days. (B) Same as (A) for Quiet Wake & REM. (C) Same as (A) for NREM. (D) Top: Log-scaled cumulative distribution function (CDF) of Active Wake bout duration in minutes. Light lines represent individual animals, dark lines represent group means, and shading represents mean ± SEM. Bottom: CDF of untransformed Active Wake bout duration. Bottom: CDFs of the non-scaled bout duration. The same analysis was performed for Quiet Wake & REM (E) and NREM (F). (E) Same as (D) for Quiet Wake/REM. (F) Same as (D) for NREM. (G) Percentage of time, number of bouts, and mean bout duration for Active Wake for each stage of the estrous cycle in female mice. Each line represents the mean value for one animal during a given estrous stage, averaged across all recorded days in that stage. Dark lines represent the mean across animals for each estrous stage; shading represents the 95% confidence interval. (H) Same as (G) for Quiet Wake/REM. (I) Same as (G) for NREM. (J) Percentage of time, number of bouts, and mean bout duration for Active Wake in 1-hour bins aligned to the light/dark cycle (Zeitgeber time, ZT), averaged across all mice and recording days. Each dot represents the mean across all mice for that bin; shading represents SEM. Dark grey shading indicates the dark phase. (K) Same as (J) for Quiet Wake & REM. (L) Same as (J) for NREM. Linear mixed-effects models were used for statistical analyses. Multiple-comparison procedures for individual analyses are reported in Table S1. n.s., not significant; \**p* < 0.05; \*\**p* < 0.01; \*\*\**p* < 0.001. Number of mice: (A–F, J–L) Male sham, n = 10; male TBI, n = 8; female sham, n = 8; female TBI, n = 11. (G–I) Numbers of mice: Female sham: proestrus, n = 4; estrus, n = 7; metestrus, n = 5; diestrus, n = 8. Female TBI: proestrus, n = 5; estrus, n = 9; metestrus, n = 5; diestrus, n = 10. For estrous-stage analyses, n represents the number of animals contributing recordings during each stage.

**Fig. S2.**
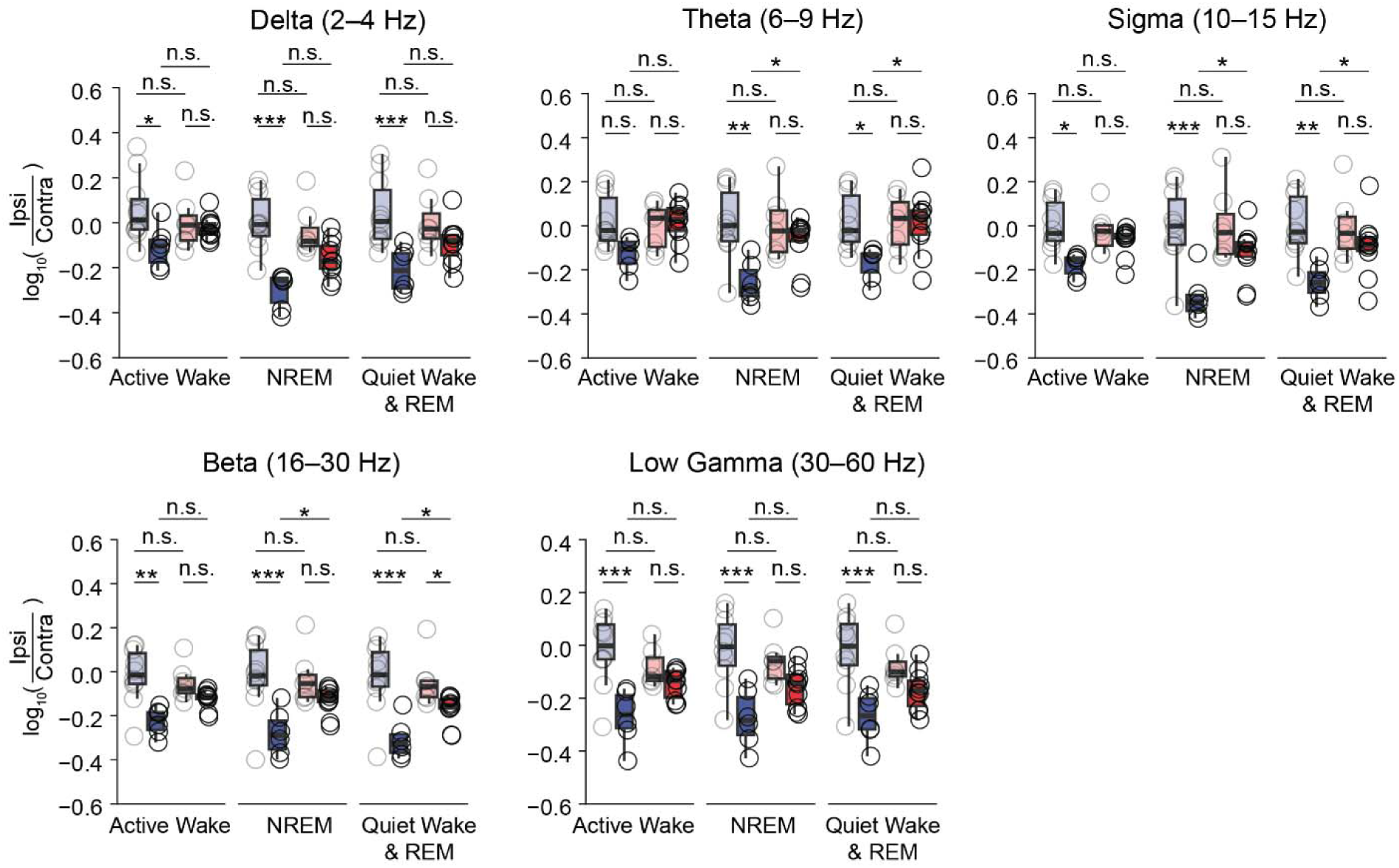
TBI broadly reduces ipsilateral relative to contralateral ECoG spectral power in males across sleep–wake states. Ratio of ipsilateral (ipsi) to contralateral (contra) ECoG power within each frequency band across sleep–wake states. Each point represents the mean value for one mouse across all bouts within the indicated sleep–wake state. The dotted line indicates an ipsilateral:contralateral ratio of 1. Number of mice: male sham, n = 10; male TBI, n = 6; female sham, n = 7; female TBI, n = 10. Linear mixed-effects models were used for statistical analyses. Multiple-comparison procedures for individual analyses are reported in Table S1. n.s., not significant; * adjusted *p* < 0.05; ** adjusted *p* < 0.01; *** adjusted *p* < 0.001.

**Fig. S3.**
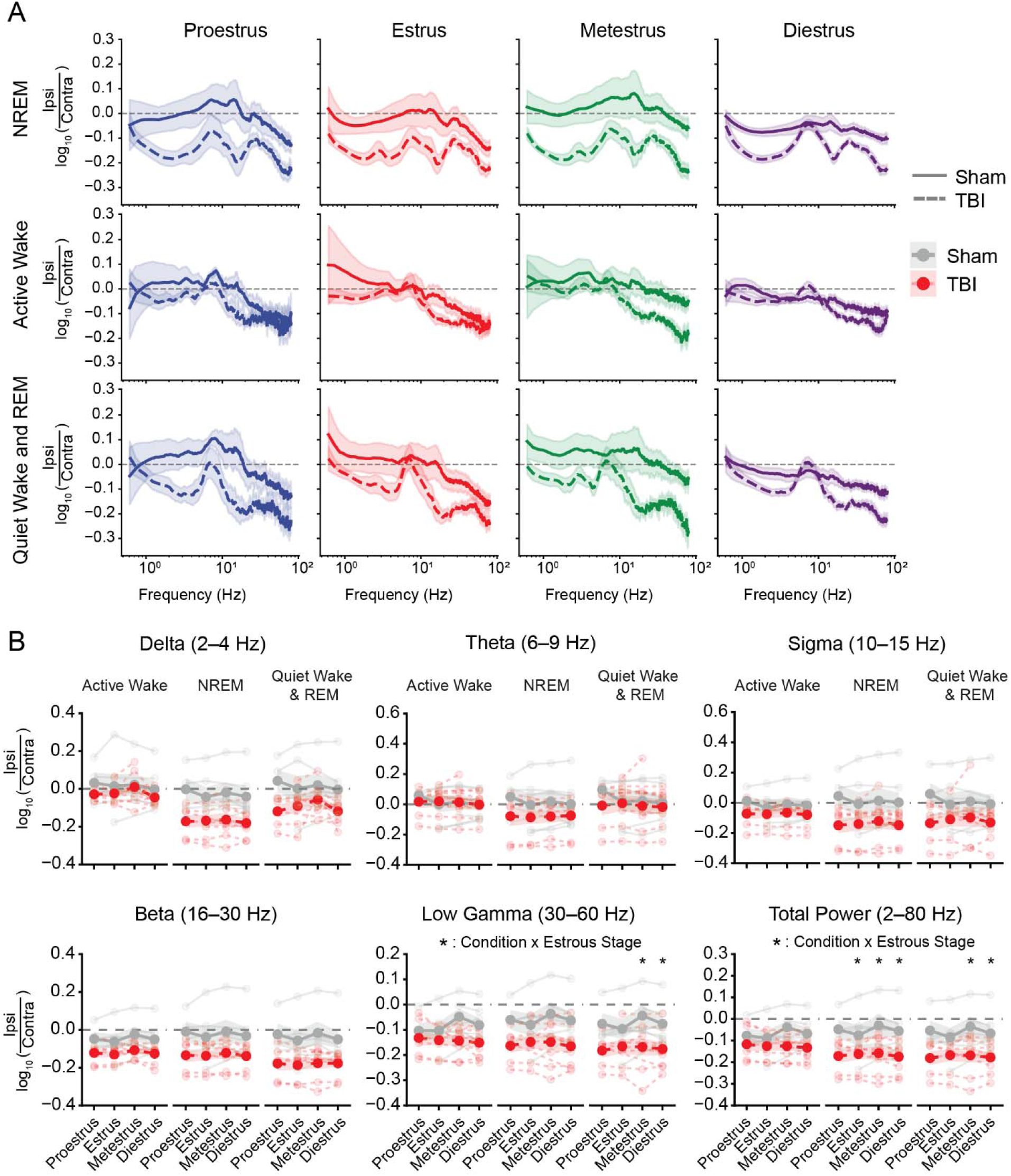
TBI-associated changes in low-gamma and total spectral power vary across the estrous cycle. (A) Ratio of ipsilateral to contralateral ECoG spectral power across frequencies for NREM, Active Wake, and Quiet Wake/REM in female mice at each estrous stage. Solid lines represent sham animals, and dashed lines represent TBI animals; shading represents SEM. (B) Ratio of ipsilateral to contralateral ECoG power within the delta (2–4 Hz), theta (6–9 Hz), sigma (10–15 Hz), beta (16–30 Hz), and low-gamma (30–60 Hz) frequency bands and for total power (2–80 Hz), shown across estrous stages for each sleep–wake state. Each light line represents the mean value for one animal during a given estrous stage, averaged across all recorded days in that stage. Dark lines represent group means. Significant TBI × estrous-stage interactions were detected for low-gamma and total power. Linear mixed-effects models were used for statistical analyses. Multiple-comparison procedures for individual analyses are reported in Table S1. * adjusted *p* < 0.05; nonsignificant post hoc comparisons are not labeled. Number of female mice: Sham: proestrus, n = 4; estrus, n = 6; metestrus, n = 5; diestrus, n = 7; TBI: proestrus, n = 6; estrus, n = 8; metestrus, n = 8; diestrus, n = 9. For estrous-stage analyses, n represents the number of animals contributing recordings during each stage.

**Fig. S4.**
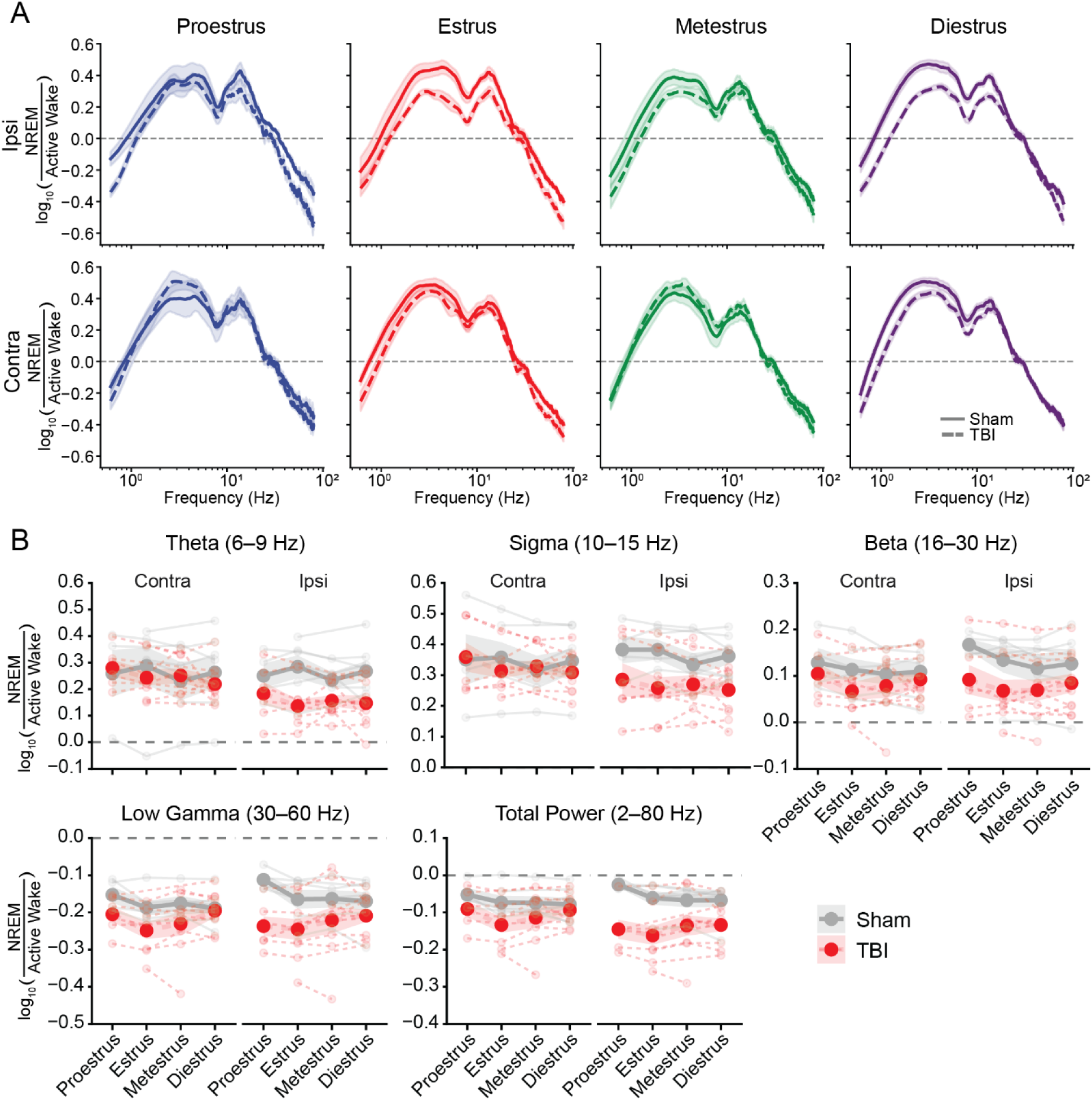
Most state-dependent spectral responses to TBI do not vary across the estrous cycle. (A) Ratio of NREM to Active Wake ECoG power across frequencies in the ipsilateral and contralateral cortex of female mice at each estrous stage. Solid lines represent sham animals and dashed lines represent TBI animals; shading represents SEM. (B) Ratio of NREM:Active Wake ECoG theta (6–9 Hz), sigma (10–15 Hz), beta (16–30 Hz), low-gamma (30–60 Hz), and total power (2–80 Hz) across the estrous cycle in the ipsilateral and contralateral cortex. Each light line represents the mean value for one animal during a given estrous stage, averaged across all recorded days in that stage. Dark lines represent group means. Note that delta and mid-gamma analyses are shown in Fig. 2D and E, respectively. Linear mixed-effects models were used for statistical analyses. Multiple-comparison procedures for individual analyses are reported in Table S1. Number of female mice: Sham: proestrus, n = 4; estrus, n = 6; metestrus, n = 5; diestrus, n = 7; TBI: proestrus, n = 6; estrus, n = 8; metestrus, n = 8; diestrus, n = 9 for the contralateral cortex and n = 10 for the ipsilateral cortex. For estrous-stage analyses, n represents the number of animals contributing recordings during each stage.

**Fig. S5.**
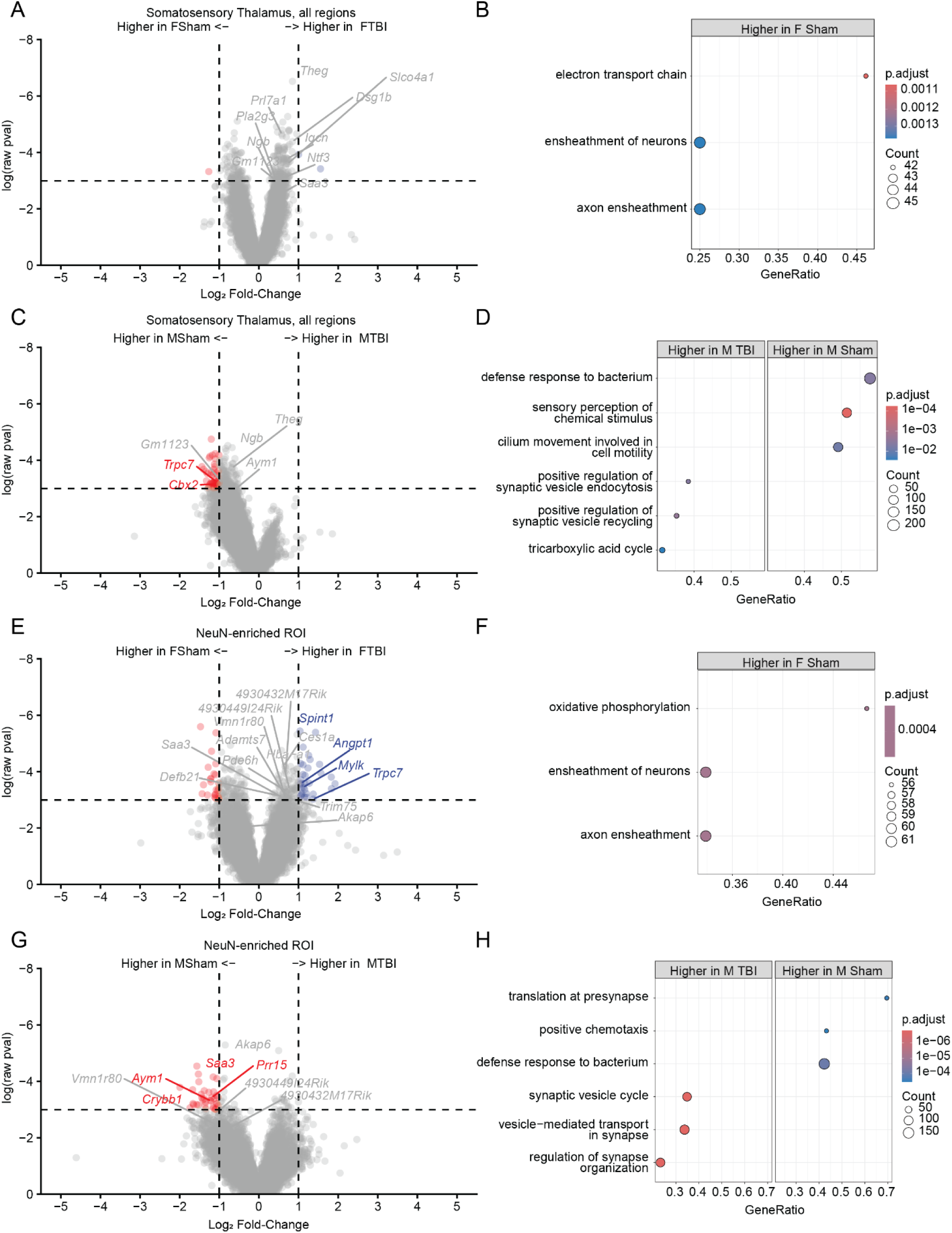
TBI produces sex-dependent transcriptional responses in the somatosensory thalamus. (A) Volcano plot comparing gene expression between female sham and female TBI mice across all ROIs in the somatosensory thalamus. Dotted lines indicate the exploratory thresholds of uncorrected *p* = 0.05 and |log□FC| = 1. Labeled genes were also identified in the female-versus-male TBI comparison shown in Fig. 7C. (B) Gene Set Enrichment Analysis of biological processes associated with differential expression between female sham and female TBI mice across all somatosensory thalamic ROIs. Significant terms were detected only among genes with higher expression in female sham mice. Dot size indicates gene count and color indicates adjusted *p* value. (C, D) Same as in (A,B) for male sham versus male TBI mice across all somatosensory thalamic ROIs. (E, F) Same analyses as in (A, B), respectively, for NeuN-enriched ROIs. (G, H) Same analyses as in (C, D), respectively, for NeuN-enriched ROIs. Linear mixed-effects models were used to calculate uncorrected *p*-values for differential-expression analyses; no individual genes passed FDR correction. Adjusted *p*-values for enrichment analyses are indicated by the color scales in (B, D, F, H). Detailed statistical results are reported in Table S1. Number of mice: Male sham, n = 3; male TBI, n = 3; female sham, n = 3; female TBI, n = 3. Number of ROIs: male sham, n = 4; male TBI, n = 5; female sham, n = 4; female TBI, n = 4. One male TBI sample contained two ROIs in order to adequately capture enough IBA1^+^ area. One set of ROIs came from a technical replicate slide as described in Fig. 7A.

**Fig. S6.**
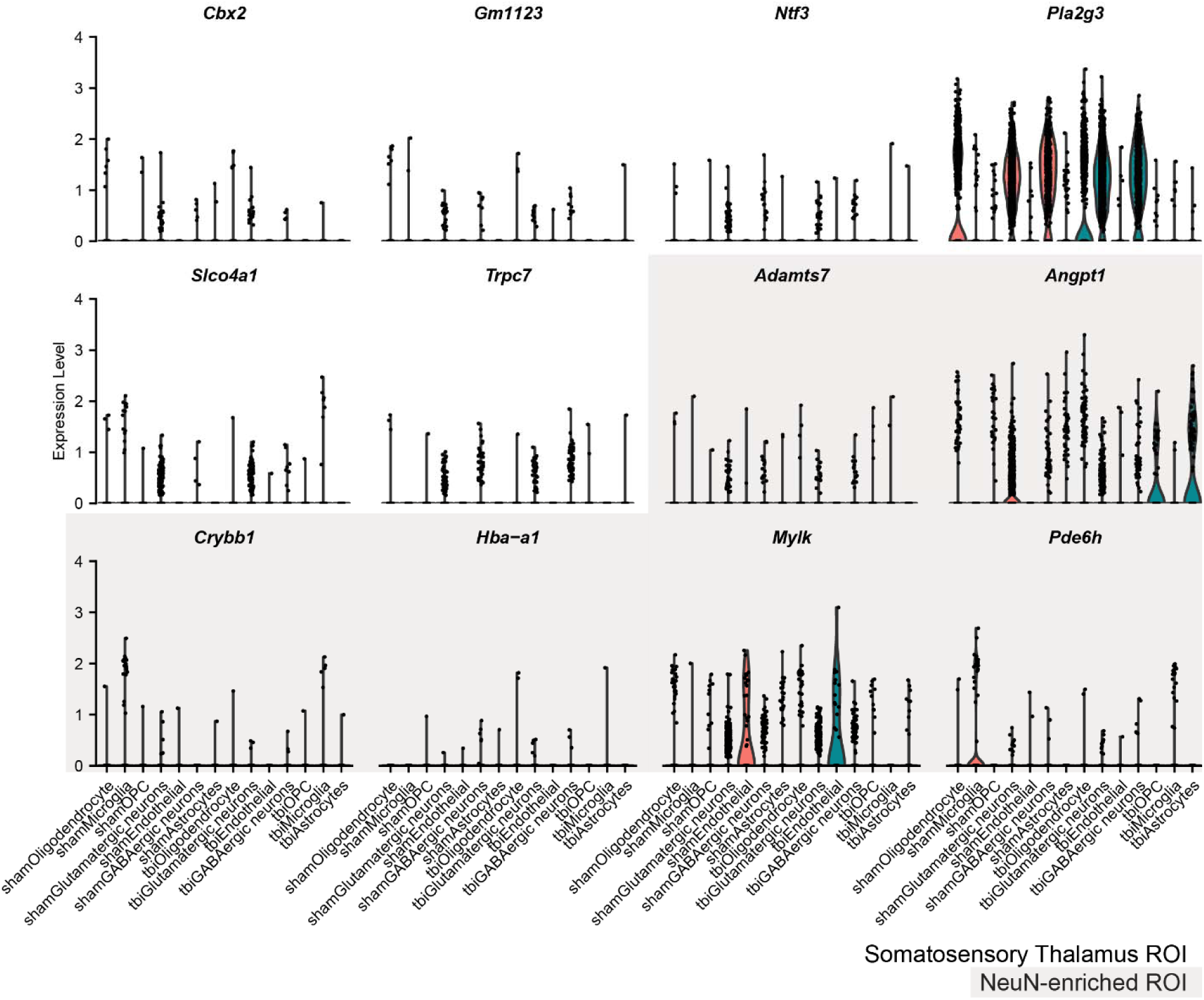
Cell-type distribution of candidate genes showing sex-dependent expression after TBI in the GeoMx dataset. Expression of the 25 candidate genes identified in the whole somatosensory thalamic region (white background) and NeuN-enriched regions (grey background) was examined across cell types in an independent, male-only snRNA-seq dataset from Holden et al. (*9*). Candidate genes were defined as genes that differed between female and male TBI mice in the GeoMx dataset (*p* < 0.05, |log□FC| > 1) and were also altered by TBI relative to sham in at least one sex (*p* < 0.05). Expression levels are shown across the indicated cell populations from the snRNA-seq dataset. Genes present in < 0.1% of cells in the snRNA-seq dataset were not plotted.

**Fig. S7.**
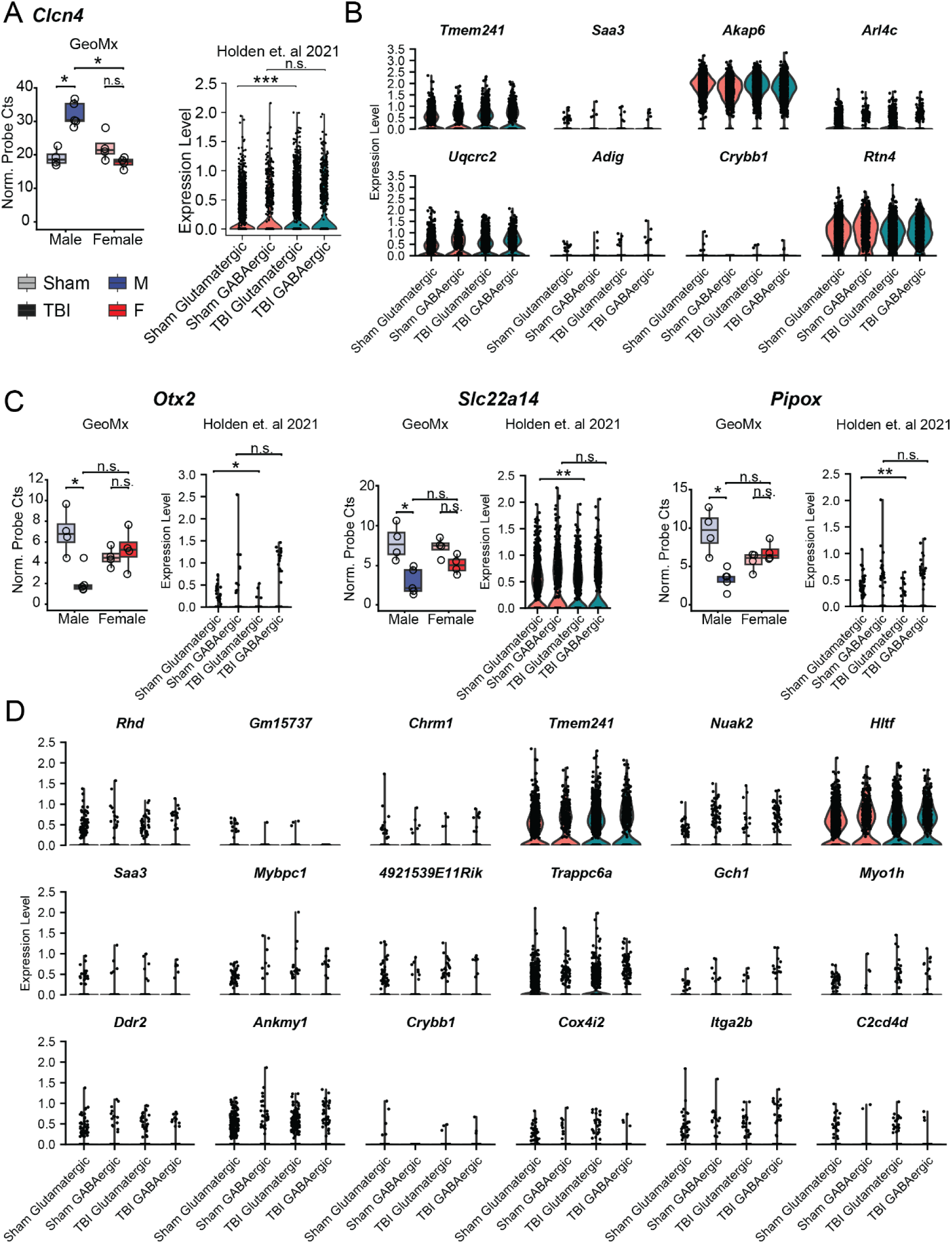
Cross-dataset comparison of candidate genes in an independent male snRNA-seq dataset. (A) *Clcn4*, which showed sex-dependent expression between TBI mice in the GeoMx dataset and was altered by TBI in males, also showed a significant TBI-associated change in an independent male snRNA-seq dataset (*9*). Left: GeoMx normalized probe counts. Right: Expression in glutamatergic and GABAergic neuronal populations from the snRNA-seq dataset. (B) Expression in the independent snRNA-seq dataset of other genes that showed sex-dependent expression between TBI mice in the GeoMx dataset and were also altered by TBI in males. Genes present in < 0.1% of cells in the snRNA-seq dataset were not plotted. (C) Same as in (A) for *Otx2, Slc22a14,* and *Pipox*, which showed large TBI-associated changes in males in the GeoMx dataset (|log□FC| > 1) but were not sex-dependently expressed after TBI. (D) Expression in the independent snRNA-seq dataset of other genes showing large TBI-associated changes in males in the GeoMx dataset (|log□FC| > 1). Genes present in < 0.1% of cells in the snRNA-seq dataset were not plotted. GeoMx probe counts were normalized using third-quartile normalization, and linear mixed-effects models were used for statistical analyses. For the snRNA-seq dataset, Wilcoxon rank-sum tests with Benjamini–Hochberg correction across tested genes were used for statistical analyses. For GeoMx comparisons, significance symbols indicate uncorrected p values. For snRNA-seq comparisons, significance symbols indicate Benjamini–Hochberg-adjusted *p*-values. n.s., not significant; * *p* < 0.05; ** *p* < 0.01; *** *p* < 0.001. Number of mice in the GeoMx dataset: Male sham, n = 3; male TBI, n = 3; female sham, n = 3; female TBI, n = 3. Number of GeoMx ROIs: male sham, n = 4; male TBI, n = 5; female sham, n = 4; female TBI, n = 4. One male TBI sample was divided into two ROIs because strong IBA1 signal prevented reliable segmentation of the somatosensory thalamus as a single ROI. One set of ROIs came from a technical replicate slide as described in Fig. 7A.

**Fig. S8.**
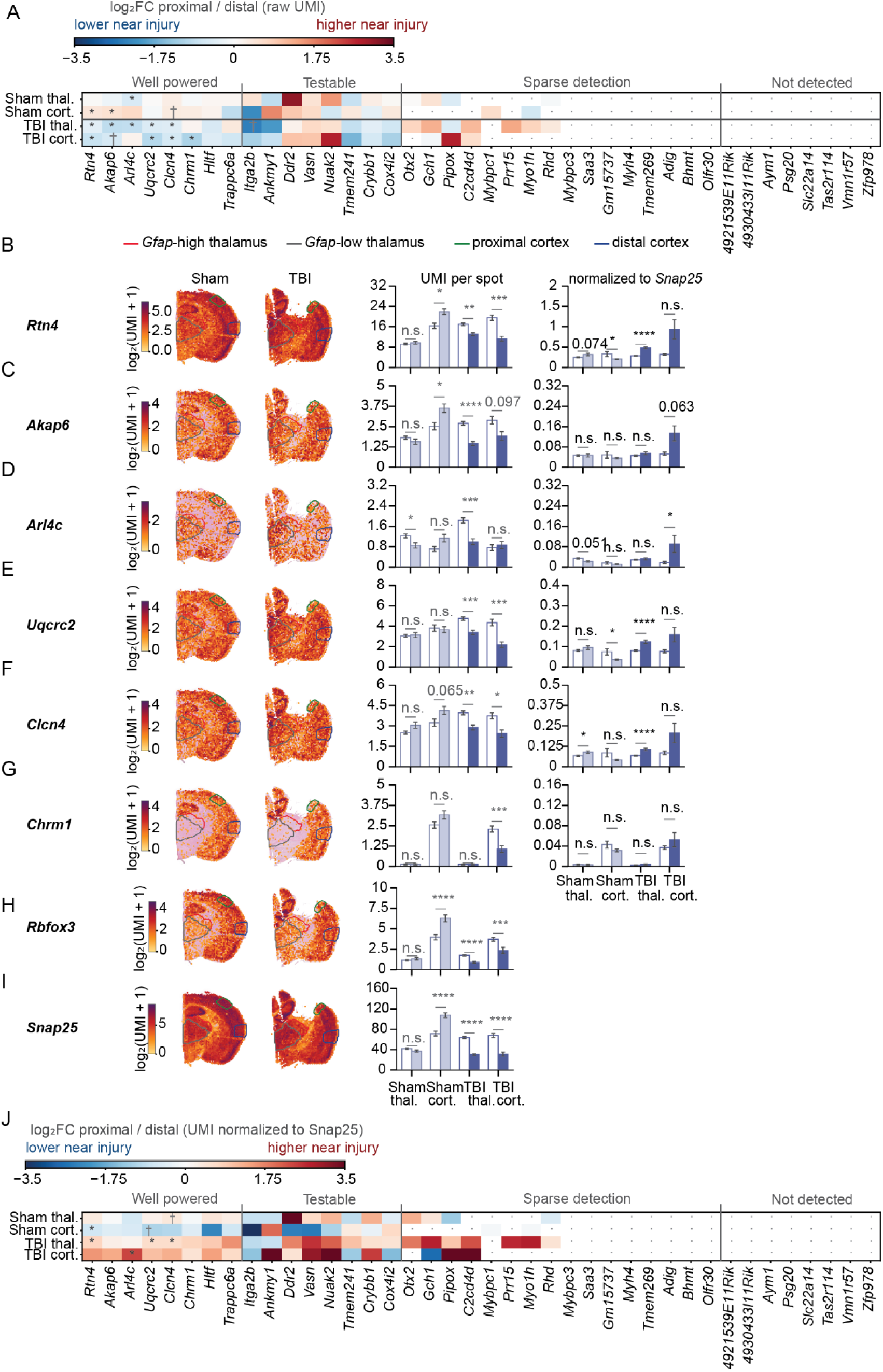
Spatial distribution of candidate gene expression in an independent Visium dataset after TBI. Re-analysis of the 10x Genomics Visium dataset from Cho et al. (*32*), comprising one sham and one TBI hemi-section from male mice 6 weeks after surgery (*n* = 1 section per condition; 2,321 sham and 2,214 TBI spots in tissue). Sections are shown with the midline on the left and cortex uppermost. Proximal/distal and *Gfap*-low/*Gfap*-high regions of interest (ROIs) were defined by Cho et al. and reused here unchanged. Thalamic ROIs were defined by unbiased k-means clustering of all spots into 10 clusters, two of which separated as *Gfap*-high and *Gfap*-low thalamic territories; cortical ROIs were defined anatomically as cortex proximal or distal to the injury. (A) Screen of all 40 candidate genes. Each cell shows the log□fold change in mean UMI per spot between paired proximal/distal or *Gfap*-high/*Gfap*-low ROIs within a single structure and section, averaged across all spots within each ROI, including spots without detected transcript (red, higher near the injury; blue, lower). Rows are grouped by condition and structure (Thal., somatosensory thalamus; Cort., cortex). Genes are ordered by the number of significant comparisons and then by their smallest adjusted *p*-value and grouped into four detection tiers: well powered (*n* = 8 genes, detected in 23–99% of ROI spots, all four ROI pairs testable), testable (*n* = 8, 3–10%, all four pairs testable), sparse detection (*n* = 16, 0–7%, zero to three pairs testable), and not detected (*n* = 8, no spot with a positive count in either section). Gray dots indicate comparisons for which no fold change was estimable, comprising 77 comparisons with too few positive spots to test and three in which one ROI contained no expressing spots. Asterisks indicate significance after multiple-comparison correction (12 of 83 testable comparisons). (B) Left: Hematoxylin-and-eosin image of each section overlaid with spots containing non-zero *Rtn4* UMI counts; color indicates log□(UMI + 1) on a scale shared between sections. Spots without detected transcripts are unmarked. Colored outlines indicate the four ROIs. Right: Mean *Rtn4* UMI per spot within each ROI, including spots without detected transcript. Open bars indicate distal ROIs and filled bars indicate proximal ROIs; light shading indicates sham and dark shading indicates TBI. Error bars represent SEM across spots. Asterisks indicate the adjusted *p*-values for the corresponding comparisons in (A). (C–G) Same as in (B) for *Akap6* (C), *Arl4c* (D), *Uqcrc2* (E), *Clcn4* (F), and *Chrm1* (G). (H, I) Spatial expression of the neuronal markers *Rbfox3* (H) and *Snap25* (I), analyzed as raw UMI counts per spot as in (B). (J) Screen of the 40 candidate genes after normalization to *Snap25*. Each cell shows the log□fold change between paired ROIs within a single structure and section following normalization of candidate-gene UMI counts to *Snap25*. Gene ordering and annotations are as in (A). *Snap25* was selected as the normalizing gene because it is measurable in nearly every spot and is among the most neuron-specific genes that are (Methods 2.4.5); normalization adjusts for the number of neurons sampled but cannot separate a change in the candidate gene from a change in *Snap25* itself. UMI, unique molecular identifier. Raw UMI counts per spot were analyzed as described previously (*32*). ROI sizes were 86 *Gfap*-high thalamic, 245 *Gfap*-low thalamic, 64 proximal cortical, and 82 distal cortical spots in the sham section, and 112, 288, 46, and 105 spots, respectively, in the TBI section. Mann–Whitney U tests were used for statistical comparisons, with Benjamini–Hochberg correction across all 83 testable comparisons. * adjusted *p* < 0.05; ** adjusted *p* < 0.01; *** adjusted *p* < 0.001; **** adjusted *p* < 0.0001. Because each condition is represented by a single section, all statistical comparisons are between ROIs within individual sections; no between-condition inference is made.

## REFERENCES

1. M. C. Dewan, A. Rattani, S. Gupta, R. E. Baticulon, Y. C. Hung, M. Punchak, A. Agrawal, A. O. Adeleye, M. G. Shrime, A. M. Rubiano, J. V. Rosenfeld, K. B. Park, Estimating the global incidence of traumatic brain injury. J Neurosurg, 1–18 (2018).

2. S. L. James, A. Theadom, R. G. Ellenbogen, M. S. Bannick, W. Montjoy-Venning, L. R. Lucchesi, N. Abbasi, R. Abdulkader, H. N. Abraha, J. C. Adsuar, M. Afarideh, S. Agrawal, A. Ahmadi, M. B. Ahmed, A. N. Aichour, I. Aichour, M. T. E. Aichour, R. O. Akinyemi, N. Akseer, F. Alahdab, A. Alebel, S. A. Alghnam, B. A. Ali, U. Alsharif, K. Altirkawi, C. L. Andrei, M. Anjomshoa, H. Ansari, M. G. Ansha, C. A. T. Antonio, S. C. Y. Appiah, F. Ariani, N. G. Asefa, S. W. Asgedom, S. Atique, A. Awasthi, B. P. Ayala Quintanilla, T. B. Ayuk, P. S. Azzopardi, H. Badali, A. Badawi, S. Balalla, A. Banstola, S. L. Barker-Collo, T. W. Bärnighausen, N. Bedi, M. Behzadifar, M. Behzadifar, B. B. Bekele, A. B. Belachew, Y. A. Belay, D. A. Bennett, I. M. Bensenor, A. Berhane, M. Beuran, A. Bhalla, S. Bhaumik, Z. A. Bhutta, B. Biadgo, M. Biffino, A. Bijani, N. Bililign, C. Birungi, S. Boufous, A. Brazinova, A. W. Brown, M. Car, R. Cárdenas, J. J. Carrero, F. Carvalho, C. A. Castañeda-Orjuela, F. Catalá-López, Y. Chaiah, A. P. Champs, J.-C. Chang, J.-Y. J. Choi, D. J. Christopher, C. Cooper, C. S. Crowe, L. Dandona, R. Dandona, A. Daryani, D. V. Davitoiu, M. G. Degefa, G. T. Demoz, K. Deribe, S. Djalalinia, H. P. Do, D. T. Doku, T. M. Drake, M. Dubey, E. Dubljanin, Z. El-Khatib, R. Ofori-Asenso, S. Eskandarieh, A. Esteghamati, S. Esteghamati, A. Faro, F. Farzadfar, M. H. Farzaei, S.-M. Fereshtehnejad, E. Fernandes, G. T. Feyissa, I. Filip, F. Fischer, T. Fukumoto, M. Ganji, F. G. Gankpe, A. K. Gebre, T. T. Gebrehiwot, K. E. Gezae, G. Gopalkrishna, A. C. Goulart, J. A. Haagsma, A. Haj-Mirzaian, A. Haj-Mirzaian, R. R. Hamadeh, S. Hamidi, J. M. Haro, H. Hassankhani, H. Y. Hassen, R. Havmoeller, C. Hawley, S. I. Hay, M. I. Hegazy, D. Hendrie, A. Henok, D. T. Hibstu, H. J. Hoffman, M. K. Hole, E. Homaie Rad, S. M. Hosseini, S. Hostiuc, G. Hu, M. A. Hussen, O. S. Ilesanmi, S. S. N. Irvani, M. Jakovljevic, S. Jayaraman, R. P. Jha, J. B. Jonas, K. M. Jones, Z. Jorjoran Shushtari, J. J. Jozwiak, M. Jürisson, A. Kabir, A. Kahsay, M. Kahssay, R. Kalani, A. Karch, A. Kasaeian, G. M. Kassa, T. D. Kassa, Z. Y. Kassa, A. P. Kengne, Y. S. Khader, M. A. Khafaie, N. Khalid, I. Khalil, E. A. Khan, M. S. Khan, Y.-H. Khang, H. Khazaie, A. T. Khoja, J. Khubchandani, A. A. Kiadaliri, D. Kim, Y.-E. Kim, A. Kisa, A. Koyanagi, K. J. Krohn, B. Kuate Defo, B. Kucuk Bicer, G. A. Kumar, M. Kumar, R. Lalloo, F. H. Lami, V. C. Lansingh, D. O. Laryea, A. Latifi, C. T. Leshargie, M. Levi, S. Li, M. L. Liben, P. A. Lotufo, R. Lunevicius, N. B. Mahotra, M. Majdan, A. Majeed, R. Malekzadeh, A.-L. Manda, M. A. Mansournia, B. B. Massenburg, K. K. V. Mate, M. M. Mehndiratta, V. Mehta, H. Meles, A. Melese, P. T. N. Memiah, W. Mendoza, G. Mengistu, A. Meretoja, T. J. Meretoja, T. Mestrovic, T. Miazgowski, T. R. Miller, G. Mini, A. Mirica, E. M. Mirrakhimov, B. Moazen, M. Mohammadi, S. Mohammed, A. H. Mokdad, M. Molokhia, L. Monasta, S. Mondello, M. Moosazadeh, G. Moradi, M. Moradi, M. Moradi-Lakeh, M. Moradinazar, S. D. Morrison, M. M. Moschos, S. M. Mousavi, S. Murthy, K. I. Musa, G. Mustafa, M. Naghavi, G. Naik, F. Najafi, V. Nangia, B. R. Nascimento, I. Negoi, T. H. Nguyen, E. Nichols, D. N. A. Ningrum, Y. L. Nirayo, P. S. Nyasulu, F. A. Ogbo, I.-H. Oh, A. Okoro, A. T. Olagunju, T. O. Olagunju, P. R. Olivares, S. S. Otstavnov, M. O. Owolabi, M. P a, S. Pakhale, A. R. Pandey, K. Pesudovs, G. D. Pinilla-Monsalve, S. Polinder, H. Poustchi, S. Prakash, M. Qorbani, A. Radfar, A. Rafay, A. Rafiei, A. Rahimi-Movaghar, V. Rahimi-Movaghar, M. Rahman, M. A. Rahman, R. K. Rai, F. Rajati, U. Ram, D. L. Rawaf, S. Rawaf, R. C. Reiner, C. Reis, A. M. N. Renzaho, S. Resnikoff, S. Rezaei, S. Rezaeian, L. Roever, L. Ronfani, G. Roshandel, N. Roy, G. M. Ruhago, B. Saddik, H. Safari, S. Safiri, M. A. Sahraian, P. Salamati, R. de F. Saldanha, A. M. Samy, J. Sanabria, J. V. Santos, M. M. M. Santric Milicevic, B. Sartorius, M. Satpathy, K. Savuon, I. J. C. Schneider, D. C. Schwebel, S. G. Sepanlou, H. Shabaninejad, M. A. A. Shaikh, M. Shams-Beyranvand, M. Sharif, M. Sharif-Alhoseini, S. M. Shariful Islam, J. She, A. Sheikh, J. Shen, K. N. Sheth, K. Shibuya, M. S. Shiferaw, M. Shigematsu, R. Shiri, I. Shiue, H. Shoman, S. Siabani, T. J. Siddiqi, J. P. Silva, D. G. A. Silveira, D. N. Sinha, M. Smith, A. M. Soares Filho, S. Sobhani, M. Soofi, J. B. Soriano, I. N. Soyiri, D. J. Stein, M. A. Stokes, M. B. Sufiyan, B. F. Sunguya, J. E. Sunshine, B. L. Sykes, C. E. I. Szoeke, R. Tabarés-Seisdedos, B. J. Te Ao, A. Tehrani-Banihashemi, M. G. Tekle, M.-H. Temsah, O. Temsah, R. Topor-Madry, M.- Tortajada-Girbés, B. X. Tran, K. B. Tran, L. Tudor Car, K. N. Ukwaja, I. Ullah, M. S. Usman, O. A. Uthman, P. R. Valdez, T. J. Vasankari, N. Venketasubramanian, F. S. Violante, F. W. S. Wagnew, Y. Waheed, Y.-P. Wang, K. G. Weldegwergs, A. Werdecker, T. Wijeratne, A. S. Winkler, G. M. A. Wyper, Y. Yano, M. Yaseri, Y. J. Yasin, P. Ye, E. M. Yimer, P. Yip, E. Yisma, N. Yonemoto, S.-J. Yoon, M. G. Yost, M. Z. Younis, M. Yousefifard, C. Yu, Z. Zaidi, S. B. Zaman, M. Zamani, Z. M. Zenebe, S. Zodpey, V. L. Feigin, T. Vos, C. J. L. Murray, Global, regional, and national burden of traumatic brain injury and spinal cord injury, 1990–2016: a systematic analysis for the Global Burden of Disease Study 2016. Lancet Neurol. 18, 56–87 (2019).

3. A. I. R. Maas, D. K. Menon, G. T. Manley, M. Abrams, C. Åkerlund, N. Andelic, M. Aries, T. Bashford, M. J. Bell, Y. G. Bodien, B. L. Brett, A. Büki, R. M. Chesnut, G. Citerio, D. Clark, B. Clasby, D. J. Cooper, E. Czeiter, M. Czosnyka, K. Dams-O’Connor, V. D. Keyser, R. Diaz-Arrastia, A. Ercole, T. A. van Essen, É. Falvey, A. R. Ferguson, A. Figaji, M. Fitzgerald, B. Foreman, D. Gantner, G. Gao, J. Giacino, B. Gravesteijn, F. Guiza, D. Gupta, M. Gurnell, J. A. Haagsma, F. M. Hammond, G. Hawryluk, P. Hutchinson, M. van der Jagt, S. Jain, S. Jain, J. Jiang, H. Kent, A. Kolias, E. J. O. Kompanje, F. Lecky, H. F. Lingsma, M. Maegele, M. Majdan, A. Markowitz, M. McCrea, G. Meyfroidt, A. Mikolić, S. Mondello, P. Mukherjee, D. Nelson, L. D. Nelson, V. Newcombe, D. Okonkwo, M. Orešič, W. Peul, D. Pisică, S. Polinder, J. Ponsford, L. Puybasset, R. Raj, C. Robba, C. Røe, J. Rosand, P. Schueler, D. J. Sharp, P. Smielewski, M. B. Stein, N. von Steinbüchel, W. Stewart, E. W. Steyerberg, N. Stocchetti, N. Temkin, O. Tenovuo, A. Theadom, I. Thomas, A. T. Espin, A. F. Turgeon, A. Unterberg, D. V. Praag, E. van Veen, J. Verheyden, T. V. Vyvere, K. K. W. Wang, E. J. A. Wiegers, W. H. Williams, L. Wilson, S. R. Wisniewski, A. Younsi, J. K. Yue, E. L. Yuh, F. A. Zeiler, M. Zeldovich, R. Zemek, Traumatic brain injury: progress and challenges in prevention, clinical care, and research. Lancet Neurol. 21, 1004–1060 (2022).

4. C. R. Baumann, E. Werth, R. Stocker, S. Ludwig, C. L. Bassetti, Sleep–wake disturbances 6 months after traumatic brain injury: a prospective study. Brain 130, 1873–1883 (2007).

5. L. L. Imbach, F. Buchele, P. O. Valko, T. Li, A. Maric, J. F. Stover, C. L. Bassetti, L. Mica, E. Werth, C. R. Baumann, Sleep-wake disorders persist 18 months after traumatic brain injury but remain underrecognized. Neurology 86, 1945–9 (2016).

6. A. Theadom, M. Cropley, P. Parmar, S. Barker-Collo, N. Starkey, K. Jones, V. L. Feigin, BIONIC Research Group, Sleep difficulties one year following mild traumatic brain injury in a population-based study. Sleep Med. 16, 926–932 (2015).

7. P. R. Kaiser, P. O. Valko, E. Werth, J. Thomann, J. Meier, R. Stocker, C. L. Bassetti, C. R. Baumann, Modafinil ameliorates excessive daytime sleepiness after traumatic brain injury. Neurology 75, 1780–1785 (2010).

8. E. M. Wickwire, J. S. Albrecht, V. F. Capaldi, S. O. Jain, R. C. Gardner, J. K. Werner, P. Mukherjee, A. B. McKeon, M. T. Smith, J. T. Giacino, L. D. Nelson, S. G. Williams, J. Collen, X. Sun, D. M. Schnyer, A. J. Markowitz, G. T. Manley, A. D. Krystal, R. Transforming, I. Clinical Knowledge in Traumatic Brain Injury, Trajectories of Insomnia in Adults After Traumatic Brain Injury. JAMA Netw Open 5, e2145310 (2022).

9. S. S. Holden, F. C. Grandi, O. Aboubakr, B. Higashikubo, F. S. Cho, A. H. Chang, A. O. Forero, A. R. Morningstar, V. Mathur, L. J. Kuhn, P. Suri, S. Sankaranarayanan, Y. Andrews-Zwilling, A. J. Tenner, A. Luthi, E. Aronica, M. R. Corces, T. Yednock, J. T. Paz, Complement factor C1q mediates sleep spindle loss and epileptic spikes after mild brain injury. Science 373, eabj2685 (2021).

10. S. S. Konduru, E. P. Wallace, J. A. Pfammatter, P. V. Rodrigues, M. V. Jones, R. K. Maganti, Sleep-wake characteristics in a mouse model of severe traumatic brain injury: Relation to posttraumatic epilepsy. Epilepsia Open 6, 181–194 (2021).

11. H. T. Korthas, B. S. Main, A. C. Harvey, R. G. Buenaventura, E. Wicker, P. A. Forcelli, M. P. Burns, The Effect of Traumatic Brain Injury on Sleep Architecture and Circadian Rhythms in Mice—A Comparison of High-Frequency Head Impact and Controlled Cortical Injury. Biology 11, 1031 (2022).

12. M. M. Lim, J. Elkind, G. Xiong, R. Galante, J. Zhu, L. Zhang, J. Lian, J. Rodin, N. N. Kuzma, A. I. Pack, A. S. Cohen, Dietary Therapy Mitigates Persistent Wake Deficits Caused by Mild Traumatic Brain Injury. Sci. Transl. Med. 5, 215ra173-215ra173 (2013).

13. M. H. Modarres, N. N. Kuzma, T. Kretzmer, A. I. Pack, M. M. Lim, EEG slow waves in traumatic brain injury: Convergent findings in mouse and man. Neurobiol. Sleep Circadian Rhythms 2, 59–70 (2017).

14. D. Necula, F. S. Cho, A. He, J. T. Paz, Secondary thalamic neuroinflammation after focal cortical stroke and traumatic injury mirrors corticothalamic functional connectivity. J. Comp. Neurol. 530, 998–1019 (2022).

15. D. W. Simon, M. McGeachy, H. Bayır, R. S. B. Clark, D. J. Loane, P. M. Kochanek, Neuroinflammation in the Evolution of Secondary Injury, Repair, and Chronic Neurodegeneration after Traumatic Brain Injury. Nat. Rev. Neurol. 13, 171–191 (2017).

16. A. Hazra, C. Macolino, M. B. Elliott, J. Chin, Delayed thalamic astrocytosis and disrupted sleep-wake patterns in a preclinical model of traumatic brain injury. J Neurosci Res 92, 1434–45 (2014).

17. S. Hébert, S. Fullum, J. Carrier, Polysomnographic and quantitative electroencephalographic correlates of subjective sleep complaints in chronic tinnitus. J. Sleep Res. 20, 38–44 (2011).

18. T. Howarth, M. Tashakori, T. Karhu, M. Rusanen, H. Pitkänen, A. Oksenberg, S. Nikkonen, Excessive daytime sleepiness is associated with relative delta frequency power among patients with mild OSA. Front. Neurol. 15, 1367860 (2024).

19. B. Lechat, C. Hirotsu, S. Appleton, M. Younes, R. J. Adams, A. Vakulin, K. Hansen, B. Zajamsek, G. Wittert, P. Catcheside, R. Heinzer, D. J. Eckert, A novel EEG marker predicts perceived sleepiness and poor sleep quality. Sleep 45, zsac051 (2022).

20. R. Paterno, H. Metheny, G. Xiong, J. Elkind, A. S. Cohen, Mild Traumatic Brain Injury Decreases Broadband Power in Area CA1. J. Neurotrauma 33, 1645–1649 (2016).

21. D. Contreras, A. Destexhe, T. J. Sejnowski, M. Steriade, Spatiotemporal Patterns of Spindle Oscillations in Cortex and Thalamus. J. Neurosci. 17, 1179–1196 (1997).

22. Y. Nir, R. J. Staba, T. Andrillon, V. V. Vyazovskiy, C. Cirelli, I. Fried, G. Tononi, Regional Slow Waves and Spindles in Human Sleep. Neuron 70, 153–169 (2011).

23. A. Osorio-Forero, R. Cardis, G. Vantomme, A. Guillaume-Gentil, G. Katsioudi, C. Devenoges, L. M. J. Fernandez, A. Lüthi, Noradrenergic circuit control of non-REM sleep substates. Curr. Biol. 31, 5009–5023.e7 (2021).

24. R. M. Jha, D. Rajasundaram, C. Sneiderman, B. T. Schlegel, C. O’Brien, Z. Xiong, K. Janesko-Feldman, R. Trivedi, V. Vagni, B. E. Zusman, J. S. Catapano, A. Eberle, S. M. Desai, A. P. Jadhav, S. Mihaljevic, M. Miller, S. Raikwar, A. Rani, J. Rulney, S. Shahjouie, I. Raphael, A. Kumar, C.-L. Phuah, E. A. Winkler, D. W. Simon, P. M. Kochanek, G. Kohanbash, A single-cell atlas deconstructs heterogeneity across multiple models in murine traumatic brain injury and identifies novel cell-specific targets. Neuron 112, 3069–3088.e4 (2024).

25. R. L. Roof, R. Duvdevani, J. W. Heyburn, D. G. Stein, Progesterone rapidly decreases brain edema: treatment delayed up to 24 hours is still effective. Exp. Neurol. 138, 246–251 (1996).

26. R. L. Roof, R. Duvdevani, D. G. Stein, Gender influences outcome of brain injury: progesterone plays a protective role. Brain Res. 607, 333–336 (1993).

27. R. L. Roof, E. D. Hall, Gender differences in acute CNS trauma and stroke: neuroprotective effects of estrogen and progesterone. J. Neurotrauma 17, 367–388 (2000).

28. J. M. Brenner, S. Ruediger, C. Wilhite, J. M. Regalado, Y. Senzai, Y. Voskobiynyk, J. T. Paz, M. Scanziani, R. Beltramo, Vision shapes neural maps of space through an ancient midbrain pathway. bioRxiv [Preprint] (2026). 10.64898/2026.05.16.725555.

29. N. Chiaruttini, C. Castoldi, L. M. Requie, C. Camarena-Delgado, B. Dal Bianco, J. Gräff, A. Seitz, B. A. Silva, ABBA+BraiAn, an integrated suite for whole-brain mapping, reveals brain-wide differences in immediate-early genes induction upon learning. Cell Rep. 44, 115876 (2025).

30. G. Yu, L.-G. Wang, Y. Han, Q.-Y. He, clusterProfiler: an R Package for Comparing Biological Themes Among Gene Clusters. OMICS J. Integr. Biol. 16, 284–287 (2012).

31. D.-D. Mao, S.-X. Li, W.-G. Hu, H. Chen, Clinical features and genetic analysis of epilepsy caused by CLCN4 gene mutation: a case report and literature review. Transl. Pediatr. 14, 1717–1725 (2025).

32. F. S. Cho, I. D. Vainchtein, Y. Voskobiynyk, A. R. Morningstar, F. Aparicio, B. Higashikubo, A. Ciesielska, D. W. M. Broekaart, J. J. Anink, E. A. van Vliet, X. Yu, B. S. Khakh, E. Aronica, A. V. Molofsky, J. T. Paz, Enhancing GAT-3 in thalamic astrocytes promotes resilience to brain injury in rodents. Sci Transl Med 14, eabj4310 (2022).

33. P. Achermann, D. J. Dijk, D. P. Brunner, A. A. Borbély, A model of human sleep homeostasis based on EEG slow-wave activity: quantitative comparison of data and simulations. Brain Res. Bull. 31, 97–113 (1993).

34. A. Borbély, The two process model of sleep regulation: Beginnings and outlook †. J. Sleep Res. 31, e13598 (2022).

35. L. M. Franke, G. T. Gitchel, R. A. Perera, R. L. Hadimani, K. L. Holloway, W. C. Walker, Randomized trial of rTMS in traumatic brain injury: improved subjective neurobehavioral symptoms and increases in EEG delta activity. Brain Inj. 36, 683–692 (2022).

36. J. S. Shaw, K. Woodard, A. Krieg, B. R. Bryant, S. Kentis, A. I. Esagoff, A. Reisch, R. M. Salas, M. E. Peters, M. J. C. Bray, The impact of traumatic brain injury on sleep and associated neuroimaging changes: A systematic review. Sleep Med. Rev. 84, 102155 (2025).

37. A. R. Adamantidis, C. Gutierrez Herrera, T. C. Gent, Oscillating circuitries in the sleeping brain. Nat. Rev. Neurosci. 20, 746–762 (2019).

38. G. Buzsáki, X.-J. Wang, Mechanisms of Gamma Oscillations. Annu. Rev. Neurosci. 35, 203–225 (2012).

39. V. S. Sohal, How Close Are We to Understanding What (if Anything) γ Oscillations Do in Cortical Circuits? J. Neurosci. 36, 10489–10495 (2016).

40. F.-C. Hsiao, P.-J. Tsai, C. W. Wu, C.-M. Yang, T. J. Lane, H.-C. Lee, L.-C. Chen, W.-K. Lee, L.-H. Lu, Y.-Z. Wu, The neurophysiological basis of the discrepancy between objective and subjective sleep during the sleep onset period: an EEG-fMRI study. Sleep 41, zsy056 (2018).

41. W. Zhao, E. J. W. Van Someren, C. Li, X. Chen, W. Gui, Y. Tian, Y. Liu, X. Lei, EEG spectral analysis in insomnia disorder: A systematic review and meta-analysis. Sleep Med. Rev. 59, 101457 (2021).

42. S. Slewa-Younan, A. M. Green, I. J. Baguley, K. L. Felmingham, A. R. Haig, E. Gordon, Is ‘gamma’ (40 Hz) synchronous activity disturbed in patients with traumatic brain injury? Clin. Neurophysiol. 113, 1640–1646 (2002).

43. E. Sanchez, H. Blais, C. Duclos, C. Arbour, S. Van Der Maren, H. El-Khatib, A.-A. Baril, F. Bernard, J. Carrier, N. Gosselin, Sleep from acute to chronic traumatic brain injury and cognitive outcomes. Sleep 45, zsac123 (2022).

44. Y. Urakami, Relationship Between Sleep Spindles and Clinical Recovery in Patients With Traumatic Brain Injury: A Simultaneous EEG and MEG Study. Clin. EEG Neurosci. 43, 39–47 (2012).

45. K. Gouveia, J. L. Hurst, Improving the practicality of using non-aversive handling methods to reduce background stress and anxiety in laboratory mice. Sci. Rep. 9, 20305 (2019).

46. F. Longordo, J. Fan, T. Steimer, C. Kopp, A. Lüthi, Do Mice Habituate to “Gentle Handling?” A Comparison of Resting Behavior, Corticosterone Levels and Synaptic Function in Handled and Undisturbed C57BL/6J Mice. Sleep 34, 679–681 (2011).

47. S. Prajapati, G. Wilbur, C. Hunley, E. Drinkard, N. Eapen, H. Mentzel, G. D. Nah, M. S. Armstrong, S. M. Murphy, R. Rowe, W. B. Hubbard, Repetitive blast TBI drives chronic sex-dependent paroxysmal motor behaviors and sleep disruption. Exp. Neurol., 116123 (2026).

48. B. Lechat, K. L. Hansen, Y. A. Melaku, A. Vakulin, G. Micic, R. J. Adams, S. Appleton, D. J. Eckert, P. Catcheside, B. Zajamsek, A Novel Electroencephalogram-derived Measure of Disrupted Delta Wave Activity during Sleep Predicts All-Cause Mortality Risk. Ann. Am. Thorac. Soc. 19, 649–658 (2022).

49. C. Arbour, S. Khoury, G. J. Lavigne, K. Gagnon, G. Poirier, J. Y. Montplaisir, J. Carrier, N. Gosselin, Are NREM sleep characteristics associated to subjective sleep complaints after mild traumatic brain injury? Sleep Med. 16, 534–539 (2015).

50. D. K. Sandsmark, M. A. Kumar, C. S. Woodward, S. E. Schmitt, S. Park, M. M. Lim, Sleep Features on Continuous Electroencephalography Predict Rehabilitation Outcomes After Severe Traumatic Brain Injury. J. Head Trauma Rehabil. 31, 101–107 (2016).

51. T. G. Rubin, M. L. Lipton, Sex Differences in Animal Models of Traumatic Brain Injury. J. Exp. Neurosci. 13, 1179069519844020 (2019).

52. D. G. Stein, Progesterone exerts neuroprotective effects after brain injury. Brain Res. Rev. 57, 386–397 (2008).

53. B. J. Allitt, V. P. A. Johnstone, K. Richards, E. B. Yan, R. Rajan, Progesterone Exacerbates Short-Term Effects of Traumatic Brain Injury on Supragranular Responses in Sensory Cortex and Over-Excites Infragranular Responses in the Long Term. J. Neurotrauma 33, 375–389 (2016).

54. H. S. Driver, D. J. Dijk, E. Werth, K. Biedermann, A. A. Borbély, Sleep and the sleep electroencephalogram across the menstrual cycle in young healthy women. J. Clin. Endocrinol. Metab. 81, 728–735 (1996).

55. Y. Ishizuka, C. P. Pollak, S. Shirakawa, T. Kakuma, K. Azumi, A. Usui, K. Shiraishi, H. Fukuzawa, T. Kariya, Sleep spindle frequency changes during the menstrual cycle. J. Sleep Res. 3, 26–29 (1994).

56. K. M. Swift, K. Keus, C. G. Echeverria, Y. Cabrera, J. Jimenez, J. Holloway, B. C. Clawson, G. R. Poe, Sex differences within sleep in gonadally intact rats. Sleep 43, zsz289 (2019).

57. A. Schroeder, M. Hudson, X. Du, Y. W. C. Wu, J. Nakamura, M. van den Buuse, N. C. Jones, R. A. Hill, Estradiol and raloxifene modulate hippocampal gamma oscillations during a spatial memory task. Psychoneuroendocrinology 78, 85–92 (2017).

58. R. L. Sumner, R. L. McMillan, A. D. Shaw, K. D. Singh, F. Sundram, S. D. Muthukumaraswamy, Peak visual gamma frequency is modified across the healthy menstrual cycle. Hum. Brain Mapp. 39, 3187–3202 (2018).

59. J. Maguire, I. Mody, Steroid Hormone Fluctuations and GABAAR Plasticity. Psychoneuroendocrinology 34, S84–S90 (2009).

60. J. L. Maguire, B. M. Stell, M. Rafizadeh, I. Mody, Ovarian cycle–linked changes in GABAA receptors mediating tonic inhibition alter seizure susceptibility and anxiety. Nat. Neurosci. 8, 797–804 (2005).

61. D. S. Reddy, Role of hormones and neurosteroids in epileptogenesis. Front. Cell. Neurosci. 7, 115 (2013).

62. D. S. Reddy, O. Gangisetty, S. Briyal, Disease-modifying activity of progesterone in the hippocampus kindling model of epileptogenesis. Neuropharmacology 59, 573–581 (2010).

63. H. E. Scharfman, N. J. MacLusky, Sex differences in the neurobiology of epilepsy: a preclinical perspective. Neurobiol. Dis. 72PB, 180–192 (2014).

64. H. E. Scharfman, N. J. MacLusky, The Influence of Gonadal Hormones on Neuronal Excitability, Seizures, and Epilepsy in the Female. Epilepsia 47, 1423–1440 (2006).

65. S. Abdulai-Saiku, S. Gupta, D. Wang, F. Marino, A. J. Moreno, Y. Huang, D. Srivastava, B. Panning, D. B. Dubal, The maternal X chromosome affects cognition and brain ageing in female mice. Nature 638, 152–159 (2025).

66. C. Lopez-Lee, L. Kodama, L. Fan, D. Zhu, J. Zhu, M. Y. Wong, P. Ye, K. Norman, N. R. Foxe, L. Ijaz, F. Yu, H. Chen, G. K. Carling, E. R. Torres, R. D. Kim, D. B. Dubal, S. A. Liddelow, S. C. Sinha, W. Luo, L. Gan, Tlr7 drives sex differences in age-and Alzheimer’s disease–related demyelination. Science 386, eadk7844 (2024).

67. M. M. McCarthy, B. M. Nugent, K. M. Lenz, Neuroimmunology and neuroepigenetics in the establishment of sex differences in the brain. Nat. Rev. Neurosci. 18, 471–484 (2017).

68. A. L. Cara, E. M. Guthman, A. Mishra, A. L. Hall, S. Yokoyama, H. N. Azadi, I. Gregg, R. M. Martinez, N. P. Sandoval, K. Aghi, K. K. Soma, S. M. Correa, A. L. Falkner, J. E. van Veen, A bifunctional optical reporter for tracking estrogen response dynamics in neurons. Neuron, doi: 10.1016/j.neuron.2026.07.031 (2026).

69. M. P. Beenhakker, J. R. Huguenard, Neurons that fire together also conspire together: is normal sleep circuitry hijacked to generate epilepsy? Neuron 62, 612–632 (2009).

70. A. Simmons, S. F. Wilferd, S. Campagnuolo, V. Pena, H. Bimonte-Nelson, J. Newbern, R. W. Sirianni, C. L. Plaisier, S. E. Stabenfeldt, Differential sex-dependent responses of circulating steroid hormones and cortical gene expression in a traumatic brain injury model. Sci. Adv. 12, eaei5171 (2026).

71. S. Villapol, D. J. Loane, M. P. Burns, Sexual dimorphism in the inflammatory response to traumatic brain injury. Glia 65, 1423–1438 (2017).

72. L. Kodama, E. Guzman, J. I. Etchegaray, Y. Li, F. A. Sayed, L. Zhou, Y. Zhou, L. Zhan, D. Le, J. C. Udeochu, C. D. Clelland, Z. Cheng, G. Yu, Q. Li, K. S. Kosik, L. Gan, Microglial microRNAs mediate sex-specific responses to tau pathology. Nat Neurosci 23, 167–171 (2020).

73. A. Villa, P. Gelosa, L. Castiglioni, M. Cimino, N. Rizzi, G. Pepe, F. Lolli, E. Marcello, L. Sironi, E. Vegeto, A. Maggi, Sex-Specific Features of Microglia from Adult Mice. Cell Rep 23, 3501–3511 (2018).

74. E. Henneman, Relation between size of neurons and their susceptibility to discharge. Science 126, 1345– 1347 (1957).

75. J. T. Paz, T. J. Davidson, E. S. Frechette, B. Delord, I. Parada, K. Peng, K. Deisseroth, J. R. Huguenard, Closed-loop optogenetic control of thalamus as a new tool to interrupt seizures after cortical injury. Nat. Neurosci. 16, 64–70 (2013).

76. J. T. Paz, C. A. Christian, I. Parada, D. A. Prince, J. R. Huguenard, Focal Cortical Infarcts Alter Intrinsic Excitability and Synaptic Excitation in the Reticular Thalamic Nucleus. J. Neurosci. 30, 5465–5479 (2010).

77. H. Fujita, K. Imura, M. Takiguchi, K. Funakoshi, Postnatal development of thalamic reticular nucleus projections to the anterior thalamic nuclei in rats. Eur. J. Histochem. EJH 66, 3370 (2022).

78. C. Song, M. Boly, E. Tagliazucchi, H. Laufs, G. Tononi, fMRI spectral signatures of sleep. Proc. Natl. Acad. Sci. U. S. A. 119, e2016732119 (2022).

79. H. Shibata, Organization of retrosplenial cortical projections to the laterodorsal thalamic nucleus in the rat. Neurosci. Res. 38, 303–311 (2000).

80. B. A. L. Perry, A. S. Mitchell, Considering the Evidence for Anterior and Laterodorsal Thalamic Nuclei as Higher Order Relays to Cortex. Front. Mol. Neurosci. 12, 167 (2019).

81. M. S. Hoseini, B. Higashikubo, F. S. Cho, A. H. Chang, A. Clemente-Perez, I. Lew, A. Ciesielska, M. P. Stryker, J. T. Paz, Gamma rhythms and visual information in mouse V1 specifically modulated by somatostatin+ neurons in reticular thalamus. eLife 10, e61437 (2021).

82. M. G. Massa, R. L. Scott, A. L. Cara, L. R. Cortes, N. P. Sandoval, J. W. Park, S. Ali, L. M. Velez, B. Tesfaye, K. Reue, J. E. van Veen, M. Seldin, S. M. Correa, Feeding Neurons Integrate Metabolic and Reproductive States in Mice. bioRxiv, 2023.01.25.525595 (2023).

83. J. B. Becker, A. P. Arnold, K. J. Berkley, J. D. Blaustein, L. A. Eckel, E. Hampson, J. P. Herman, S. Marts, W. Sadee, M. Steiner, J. Taylor, E. Young, Strategies and methods for research on sex differences in brain and behavior. Endocrinology 146, 1650–1673 (2005).

84. M. J. Prerau, R. E. Brown, M. T. Bianchi, J. M. Ellenbogen, P. L. Purdon, Sleep Neurophysiological Dynamics Through the Lens of Multitaper Spectral Analysis. Physiology 32, 60–92 (2017).

85. L. M. J. Fernandez, A. Lüthi, Sleep Spindles: Mechanisms and Functions. Physiol. Rev. 100, 805–868 (2020).

86. L. M. Fernandez, G. Vantomme, A. Osorio-Forero, R. Cardis, E. Béard, A. Lüthi, Thalamic reticular control of local sleep in mouse sensory cortex. eLife 7, e39111 (2018).

87. C. R. Merritt, G. T. Ong, S. E. Church, K. Barker, P. Danaher, G. Geiss, M. Hoang, J. Jung, Y. Liang, J. McKay-Fleisch, K. Nguyen, Z. Norgaard, K. Sorg, I. Sprague, C. Warren, S. Warren, P. J. Webster, Z. Zhou, D. R. Zollinger, D. L. Dunaway, G. B. Mills, J. M. Beechem, Multiplex digital spatial profiling of proteins and RNA in fixed tissue. Nat. Biotechnol. 38, 586–599 (2020).

